# Sonic Hedgehog Activates Prostaglandin Signaling to Stabilize Primary Cilium Length

**DOI:** 10.1101/2022.05.06.490951

**Authors:** Shariq S. Ansari, Miriam E. Dillard, Yan Zhang, Mary Ashley Austria, Naoko Boatwright, Elaine L. Shelton, Amanda Johnson, Brandon M. Young, Zoran Rankovic, John D. Schuetz, Camenzind G. Robinson, Stacey K. Ogden

## Abstract

Sonic Hedgehog (SHH) signaling is an essential driver of embryonic patterning that, when corrupted, leads to developmental disorders and cancers. SHH effector responses are organized through nonmotile primary cilia that grow and retract with the cell cycle and in response to distinct extracellular cues. Destabilization of primary cilium length corrupts SHH pathway regulation, which places significant pressure on SHH to maintain ciliary architecture. Herein, we investigate whether SHH signaling promotes ciliary length control. We reveal a signal crosstalk circuit induced by SHH activation of Phospholipase A2 (cPLA_2_) that drives ciliary EP_4_ signaling to stabilize primary cilium length. We demonstrate that chemical or genetic blockade of SHH-EP_4_ crosstalk leads to destabilized primary cilium cyclic AMP (cAMP) control, reduced ciliary length, and attenuated SHH pathway induction. Accordingly, we find that *Ep_4_ ^-/-^* mice display shortened neuroepithelial primary cilia and altered SHH-dependent neuronal cell fate specification. Thus, SHH initiates a signaling crosstalk circuit that maintains primary cilium length for a robust downstream signaling response.

## INTRODUCTION

During embryonic patterning, Sonic Hedgehog (SHH) signaling proteins provide instructional cues that guide cell fate decisions to organize tissues. Disruption of SHH pathway activation leads to developmental disorders and, consistent with regulated signaling being important for tissue homeostasis post-development, aberrant SHH activity drives cancer (Barakat et al., 2010). As such, sophisticated regulatory mechanisms are in place to ensure appropriate pathway control in both basal and activated signaling states. Tight regulation of pathway activity requires localization of SHH pathway components to a specialized sensory organelle called the primary cilium (Caspary et al., 2007; Huangfu and Anderson, 2006). Each cell possesses a single primary cilium that is anchored to the basal body and allows for organization of numerous signal transduction cascades to instruct diverse cellular responses (Anvarian et al., 2019; Wheway et al., 2018). Despite the membrane of the primary cilium being contiguous with the plasma membrane, it maintains a distinct membrane lipid composition through action of lipid metabolic enzymes that localize near the ciliary base (Arensdorf et al., 2017; Findakly et al., 2021; Garcia et al., 2018; Raleigh et al., 2018). Further, a cytoplasmic diffusion barrier controls entry of soluble proteins into the ciliary body, ensuring that primary cilia offer protected environments for interpretation and transduction of myriad extracellular signals (Garcia et al., 2018; Shi et al., 2017; Ye et al., 2013).

The SHH receptor Patched (PTCH), signal transducer Smoothened (SMO), and GLI2/GLI3 transcriptional effectors all function at or cycle through primary cilia in ligand-regulated manners (Arensdorf et al., 2016; Petrov et al., 2017). In the absence of SHH, SMO retention in primary cilia is restricted by PTCH-mediated depletion of SMO-activating sterols from ciliary membrane (Kinnebrew et al., 2019; Kinnebrew et al., 2021; Kong et al., 2019). In this off state, phosphorylation of GLI2 and GLI3 transcription factors by cyclic AMP (cAMP)-dependent protein kinase (PKA) in primary cilia tags them for partial degradation to remove their transcriptional activation domains (Bangs and Anderson, 2017; Haycraft et al., 2005; Tuson et al., 2011). SHH binding to PTCH attenuates PTCH-mediated ciliary sterol elimination, which leads to SMO sterol binding and retention in primary cilia where it signals to block PKA-promoted GLI truncation.

This allows for accumulation of full-length GLI2/3 proteins that are subsequently activated for nuclear translocation and target gene induction (Niewiadomski et al., 2014; Umberger and Ogden, 2021).

Current models suggest dependency of SHH signaling on primary cilia evolved due to the specialized lipid composition of ciliary membrane allowing for tight control over availability of SMO-activating sterols (Kinnebrew et al., 2019; Kinnebrew et al., 2021; Kong et al., 2019; Raleigh et al., 2018). Further, the small size of primary cilia allows for rapid modulation of second messengers such as cAMP (Hansen et al., 2020; Truong et al., 2021). Consistent with this functionality and the key role of PKA in regulating GLI that cycles through primary cilia, SHH signal output is highly sensitive to changes in ciliary cAMP concentration (Mukhopadhyay et al., 2013; Truong et al., 2021; Vuolo et al., 2015). Pathway activation lowers primary cilium cAMP to reduce PKA activity by stimulating ciliary exit of Gα_s_-coupled GPR161 (Mukhopadhyay et al., 2013). In addition, SMO can activate heterotrimeric G proteins of the Gα_i_ family to inhibit adenylyl cyclase (AC), suggesting that high-level activation of SHH signaling can lead to sustained reduction in primary cilium cAMP (Barzi et al., 2011; Ogden et al., 2008; Riobo and Manning, 2007).

Paradoxically, a growing body of evidence suppxsxsxsxsxorts that ciliary length stability requires high ciliary cAMP. Studies in zebrafish demonstrate that Multidrug Resistance Protein 4 Transporter (ABCC4), which exports the prostaglandin signaling molecule PGE_2_, is required for motile cilium elongation. Ciliary length is regulated by PGE_2_ activation of the ciliary G protein coupled receptor (GPCR) E-type prostanoid receptor 4 (EP_4_) (Jin et al., 2014; Marra et al., 2019). Active EP_4_ stimulates Gα_s_ heterotrimeric G proteins, which raise intraciliary cAMP to promote anterograde intraflagellar transport (IFT) for motile cilium length maintenance (Besschetnova et al., 2010; Hansen et al., 2020; Hansen et al., 2022; Jin et al., 2014). Primary cilium length control likely occurs through similar mechanisms because reduced ABCC4 activity in cultured fibroblasts leads to primary cilium shortening and increasing ciliary cAMP leads to primary cilium elongation (Besschetnova et al., 2009; Jin et al., 2014). This raises the possibility that ciliary cAMP reduction in response to high-level activation of SHH signaling could slow IFT, shorten primary cilia, and blunt the downstream signal response. The fail-safes that prevent ciliary shortening in SHH-stimulated cells have not been described.

We previously reported that SHH activates cytosolic phospholipase A2α (cPLA_2_) through SMO activation of Gβγ to produce arachidonic acid. Arachidonic acid drives a feed-forward signal that enhances SMO ciliary enrichment such that inhibition of cPLA_2_ attenuates agonist-induced SMO ciliary accumulation and blunts signaling to GLI transcription factors (Arensdorf et al., 2017).

Notably, arachidonic acid can be metabolized to generate the EP_4_ agonist PGE_2_. This suggests that, in addition to enhancing SMO ciliary enrichment, SHH-mediated production of arachidonic acid may connect the SHH pathway with PGE_2_/EP_4_-regulated primary cilium length control.

Herein, we investigate this hypothesis and reveal a crosstalk circuit between SHH and prostaglandin signaling pathways. We report that SHH-stimulated production of arachidonic acid drives ciliary length homeostasis through production of PGE_2_ and activation of EP_4_. This provides a mechanism by which ciliary cAMP levels are stabilized in SHH-stimulated cells to sustain anterograde IFT and prevent primary cilium shortening. We show that genetic ablation or small molecule inhibition of SHH-to-EP_4_ crosstalk destabilizes primary cilium cAMP concentration, slows IFT, shortens primary cilia, reduces SMO ciliary enrichment, and attenuates SMO signaling to GLI. Consistent with EP_4_ signaling being crucial for regulated SHH pathway activation *in vivo*, we find that mice lacking EP_4_ have shortened primary cilia and exhibit neural tube patterning defects indicative of compromised SHH signaling. As such, we propose the SHH pathway evolved to link with prostaglandin signaling through cPLA_2_ activation to stabilize primary cilium length for optimal function of the organelle from which it instructs downstream signaling.

## RESULTS

### cPLA_2_ contributes to ciliary length control downstream of SHH

To determine whether SHH activation of cPLA_2_ leads to secretion of PGE_2_ that regulates primary cilium morphology, we turned to murine kidney inner medullary collecting duct (IMCD3) cells. IMCD3 cells are SHH responsive, produce long primary cilia, and provide an established model for examination of ciliary biology (Breslow et al., 2018; May et al., 2021; Ott and Lippincott-Schwartz, 2012; Zhou et al., 2014). IMCD3 cells were stimulated with control conditioned media or conditioned media containing the amino-terminal signaling domain of SHH for ∼24 hours, and then media was collected to test for accumulation of PGE_2_. Enzyme-linked immunosorbent assays (ELISA) of conditioned media from control and ligand-treated IMCD3 cells showed that SHH stimulated release of PGE_2_ and that release was blocked by pretreatment with the inverse SMO agonist cyclopamine. SHH-stimulated PGE_2_ secretion was also blocked by pre-treatment with the specific cPLA_2_ inhibitor giripladib (GIRI) (Figure 1A) (Arensdorf et al., 2017; Duvernay et al., 2015). To determine whether PGE_2_ secretion occurred in an ABCC4-dependent manner, we measured release from control or *Abcc4* knockout NIH 3T3 cells following SHH stimulation. Similar to what was observed with IMCD3 cells, SHH stimulation of control NIH-3T3 cells induced secretion of PGE_2_ into cell culture medium. This secretion was blocked by inhibiting SMO with cyclopamine or by inhibiting the cyclooxygenase (COX) responsible for initiating arachidonic acid to PGE_2_ conversion (Figure 1A’, Coxib).

**Figure 1:**
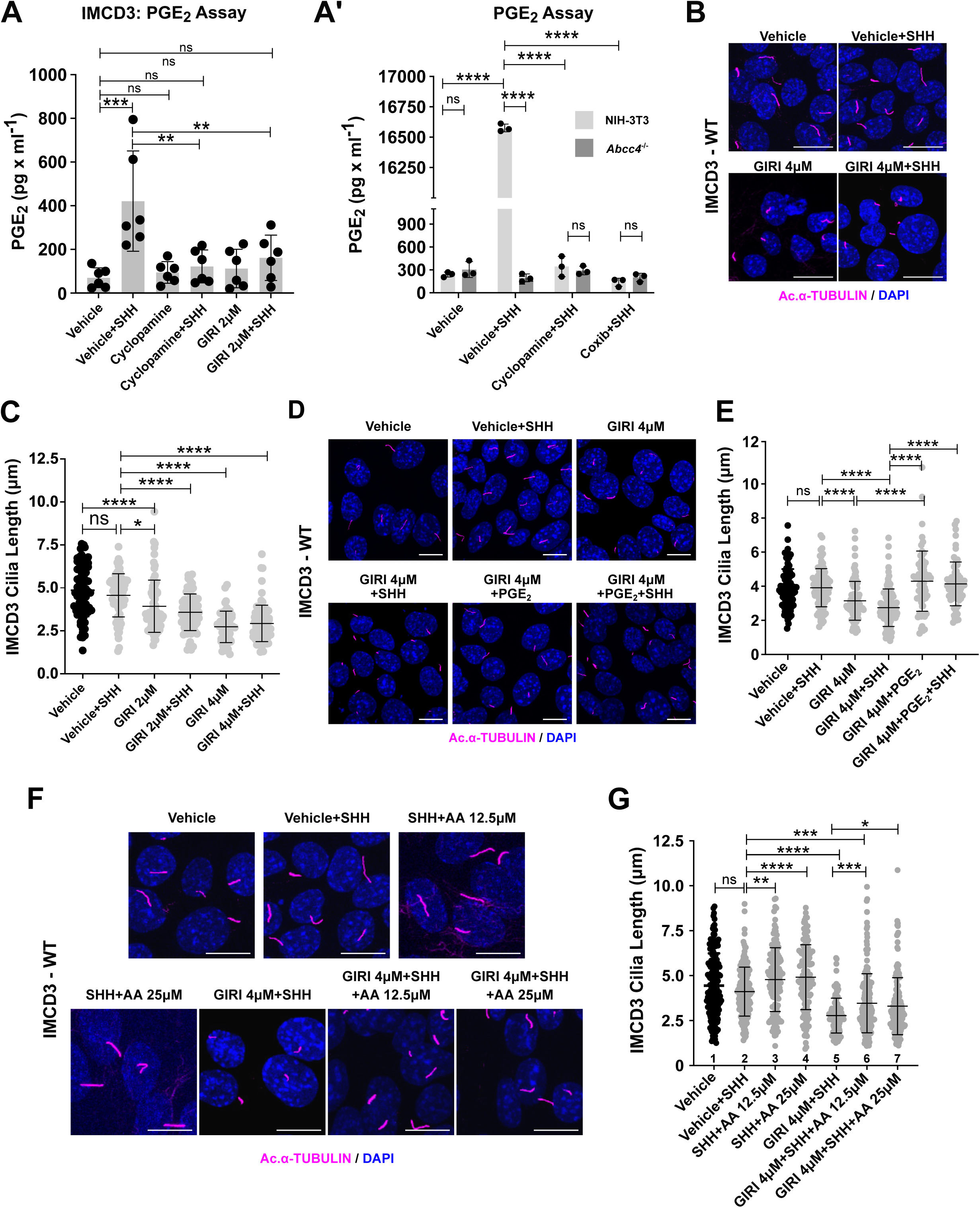
PGE_2_ generated downstream of SHH contributes to ciliary length control. **(A-A’)**. PGE_2_ is secreted downstream of SMO and cPLA_2_ activation. PGE_2_ was measured in IMCD3 **(A)** or WT and *Abcc4^-/-^*NIH-3T3 **(A’)** cell culture media by ELISA. **(A).** Cells were pretreated with vehicle control (DMSO), the inverse SMO agonist cyclopamine (10 µM), or cPLA_2_ inhibitor giripladib (GIRI, 2 µM), and then exposed to SHH. PGE_2_ was measured in cell culture supernatant ∼36 hours post treatment. The experiment was repeated twice with 3 biological replicates per experiment. Pooled data are shown. **(A’).** Cells were pretreated with vehicle (DMSO), cyclopamine (10 µM), or COX2 inhibitor Celecoxib (Coxib,10 µM) and then stimulated with control or SHH conditioned media. PGE_2_ was measured in the supernatant ∼36 hours after treatment. The experiment was performed twice with 3 biological replicates per experiment. A representative experiment is shown. **(B-C).** cPLA_2_ inhibition shortens primary cilia. **(B)**. IMCD3 cells were treated with SHH conditioned media in the presence of GIRI (4 µM), or vehicle control (DMSO). The ciliary axoneme is marked by acetylated α-tubulin (magenta). DAPI (blue) marks the nucleus. Scale bar = 10 μm. **(C)**. Lengths of primary cilia were quantified in IMCD3 cells treated with SHH conditioned media in the absence or presence of GIRI. Average primary cilium length was determined by measuring cilia of ≥100 cells/condition across three independent experiments. **(D-E).** PGE_2_ supplementation rescues ciliary length in GIRI-treated IMCD3 cells. Cells were treated with 4 µM GIRI in the presence of PGE_2_ media supplementation (40 µM) or vehicle control. Scale bar = 10 μm. Ciliary length quantification is shown in **E. (F-G).** Arachidonic acid supplementation rescues primary cilium length following cPLA_2_ inhibition. IMCD3 cells were treated with SHH conditioned media in the presence of vehicle or 4 µM GIRI in media supplemented with increasing concentrations of arachidonic acid (12.5 µM or 25 µM). Scale bar = 10 μm. Ciliary length quantification is shown in **G**. For all experiments, statistical significance was calculated using one-way ANOVA. A p value of less than 0.05 was considered statistically significant with significance indicated as follows: *<0.05, **<0.01, ***<0.001, ****<0.0001, and ns, p > 0.05. Data are represented as mean ± SD.

Consistent with ABCC4 secreting PGE_2_ that is generated in response to SHH pathway activation, *Abcc4* knockout NIH-3T3 cells failed to release prostaglandin in response to SHH treatment (Figure 1A’, light gray vs. dark gray). As such, arachidonic acid produced downstream of SHH-stimulated cPLA_2_ provides a substrate for PGE_2_ production and secretion.

Having established that SHH-mediated cPLA_2_ activation stimulates PGE_2_ release from IMCD3 and NIH-3T3 cells, we next evaluated the effect of reducing cPLA_2_-generated arachidonic acid on ciliary length. We treated IMCD3 cells with vehicle or GIRI in the absence and presence of SHH, and measured lengths of acetylated α-tubulin-marked primary cilia. We generated maximum projections of Z stack images, and then traced the length of each acetylated α-tubulin marked cilium from base to the tip using the line profile tool in LAS X software. GIRI treatment of IMCD3 cells shortened primary cilia significantly, independent of SHH stimulation status.

Primary cilium length averaged ∼5 μm in the absence of GIRI and shortened by approximately 40% to ∼3 μm at the highest GIRI concentration used (Figure 1B-C). Ciliary lengths of GIRI treated cells were rescued by addition of supplemental PGE_2_ or its precursor arachidonic acid (Figure 1D-G). Notably, supplemental arachidonic acid augmented ciliary length in SHH stimulated cells in the absence of GIRI, further supporting a link between increased arachidonic acid availability and primary cilium elongation (Figure 1G, column 2 vs. 3-4). Taken together, these results support that ciliary shortening observed in GIRI-treated cells was a specific effect of compromised arachidonic acid-to-PGE_2_ metabolism.

To determine whether the ciliary morphology changes that occurred following cPLA_2_ inhibition correlated with reduced SMO ciliary accumulation in IMCD3 cells, we stimulated with SHH conditioned media in the absence or presence of GIRI and quantified SMO ciliary signal intensity (Supplementary Figure 1A-E’). Consistent with what we previously reported for NIH 3T3 cells (Arensdorf et al., 2017), cPLA_2_ inhibition disrupted SMO ciliary retention in SHH treated cells (Supplementary Figure 1A-E, red). SHH-stimulated ciliary exit of GPR161 (magenta) and retention of the primary cilium resident protein ARL13B (green) were unaffected by GIRI treatment (Supplementary Figure 1A-D, E’), suggesting specificity in the ciliary functions affected by cPLA_2_ inhibition.

### EP_4_ activation promotes ciliary length homeostasis for SHH signaling to GLI

Based on established connections between prostaglandin signaling and motile cilia elongation (Jin et al., 2014; Marra et al., 2019), we hypothesized PGE_2_ produced downstream of SHH signals through ciliary EP_4_ to influence primary cilium length. We therefore sought to determine whether EP_4_ localized to primary cilia of IMCD3 cells. EP_4_ was evident in primary cilia of both IMCD3 cells and control mouse embryonic fibroblasts (MEF) and was undetectable in cilia of *Ep ^-/-^* MEFs. These results support specificity of the EP_4_ antibody signal and demonstrate that EP_4_ localizes to primary cilia in both MEFs and IMCD3 cells (Figure 2A-C’).

**Figure 2:**
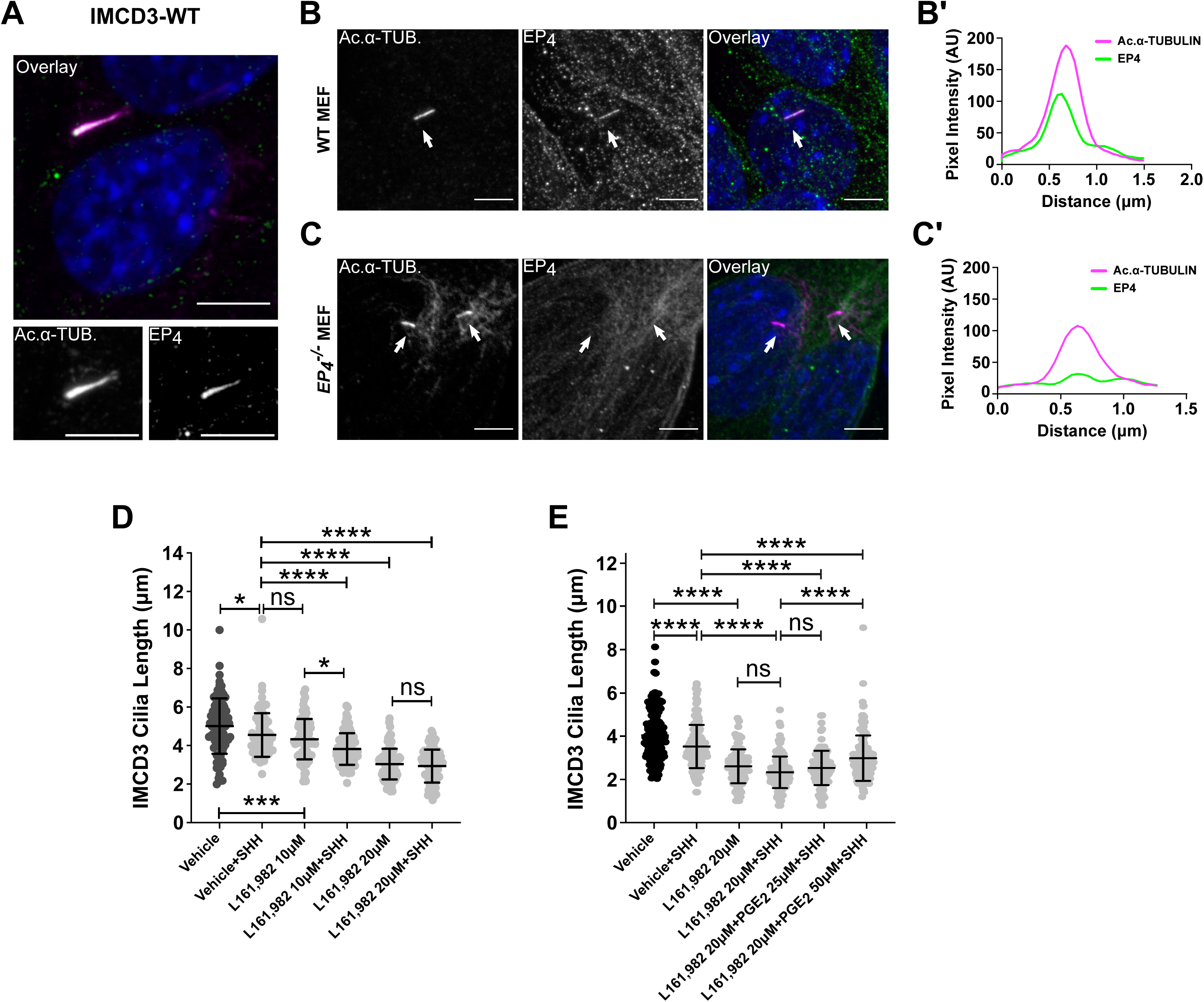
EP_4_ is required for primary cilium length homeostasis. **(A).** EP_4_ localizes to primary cilia of IMCD3 cells. Endogenous EP_4_ is shown in green and acetylated α-tubulin (magenta) marks the primary cilium. Scale bar = 5 µm. **(B-C’).** EP_4_ localizes to primary cilia in wild type MEFs. Ciliary EP_4_ signal is absent in *Ep_4_ ^-/-^*cells. Line scans indicate the degree of localization between EP_4_ (green) and acetylated α-tubulin (magenta) within the primary cilium in control **(B’)** and *Ep ^-/-^*MEFs **(C’)**. Intensity profiles are presented as arbitrary units (AU). Scale bars = 5 μm. **(D).** EP_4_ inhibition shortens primary cilia in IMCD3 cells. Cells were pretreated with vehicle control (DMSO) or EP_4_ inhibitor L161,982 (10 µM or 20 µM) for two hours prior to addition of SHH. **(E).** PGE_2_ partially rescues primary cilia length following EP_4_ inhibition. IMCD3 cells were treated with SHH conditioned media in the presence of vehicle or 20 µM L161,982 in media supplemented with increasing concentrations of PGE_2_ (25 or 50 μM). At least 150 cilia/condition were measured over three experiments. Statistical significance was calculated using one-way ANOVA. For all experiments, a p value of less than 0.05 was considered statistically significant with significance indicated as follows: *<0.05, **<0.01, ***<0.001, ****<0.0001, and ns, p > 0.05. Data are represented as mean ± SD.

To test for EP_4_ contributions to primary cilium length maintenance, average ciliary lengths were calculated in control and SHH-treated IMCD3 cells exposed to vehicle or the high-affinity EP_4_ inhibitor, L161,982 (Cherukuri et al., 2007). Primary cilia of IMCD3 cells treated with increasing concentrations of L161,982 shortened in a dose-dependent manner in the absence of SHH. Notably, ciliary shortening of cells treated with low-dose L161,982 was significantly enhanced by SHH treatment (Figure 2D), demonstrating that an active SHH pathway disrupts primary cilium length control if EP_4_ activity is compromised. Supplementing culture media of L161,982-treated cells with high concentrations of the EP_4_ ligand PGE_2_ partially rescued ciliary length of SHH stimulated cells, supporting that primary cilium shortening following L161,982 treatment was a specific effect of on-target inhibitor activity toward EP_4_ (Figure 2E).

Having determined that EP_4_ contributes to ciliary length maintenance in SHH-stimulated cells, we next wanted to understand how EP_4_ inhibition affected SHH signaling output. Thus, SHH stimulated SMO ciliary signal intensity was quantified in control, GIRI-treated, and L161,982 treated IMCD3 cells. Similar to what we observed following cPLA_2_ inhibition with GIRI, inhibition of EP_4_ with L161,982 attenuated ciliary accumulation of endogenous SMO following SHH exposure (Figure 3A-B and Supplementary Figure 1). Ciliary shortening and reduced SMO accumulation relative to control were also observed in *Ep ^-/-^* MEFs (Figure 3C-E), further supporting on-target activity of L161,982 to EP_4_. To determine whether reduced SMO ciliary enrichment following EP_4_ inhibition compromised activation of GLI transcriptional effectors, qPCR analyses of SHH transcriptional targets *Gli1* and *Ptch1* were performed in IMCD3 and NIH-3T3 cells. Inhibition of either cPLA_2_ with GIRI or EP_4_ with L161,982 significantly reduced the ability of SHH to activate a transcriptional response (Figure 3F-F’). Thus, compromised PGE_2_/EP_4_ signaling shortens cilia and attenuates SHH pathway activity.

**Figure 3:**
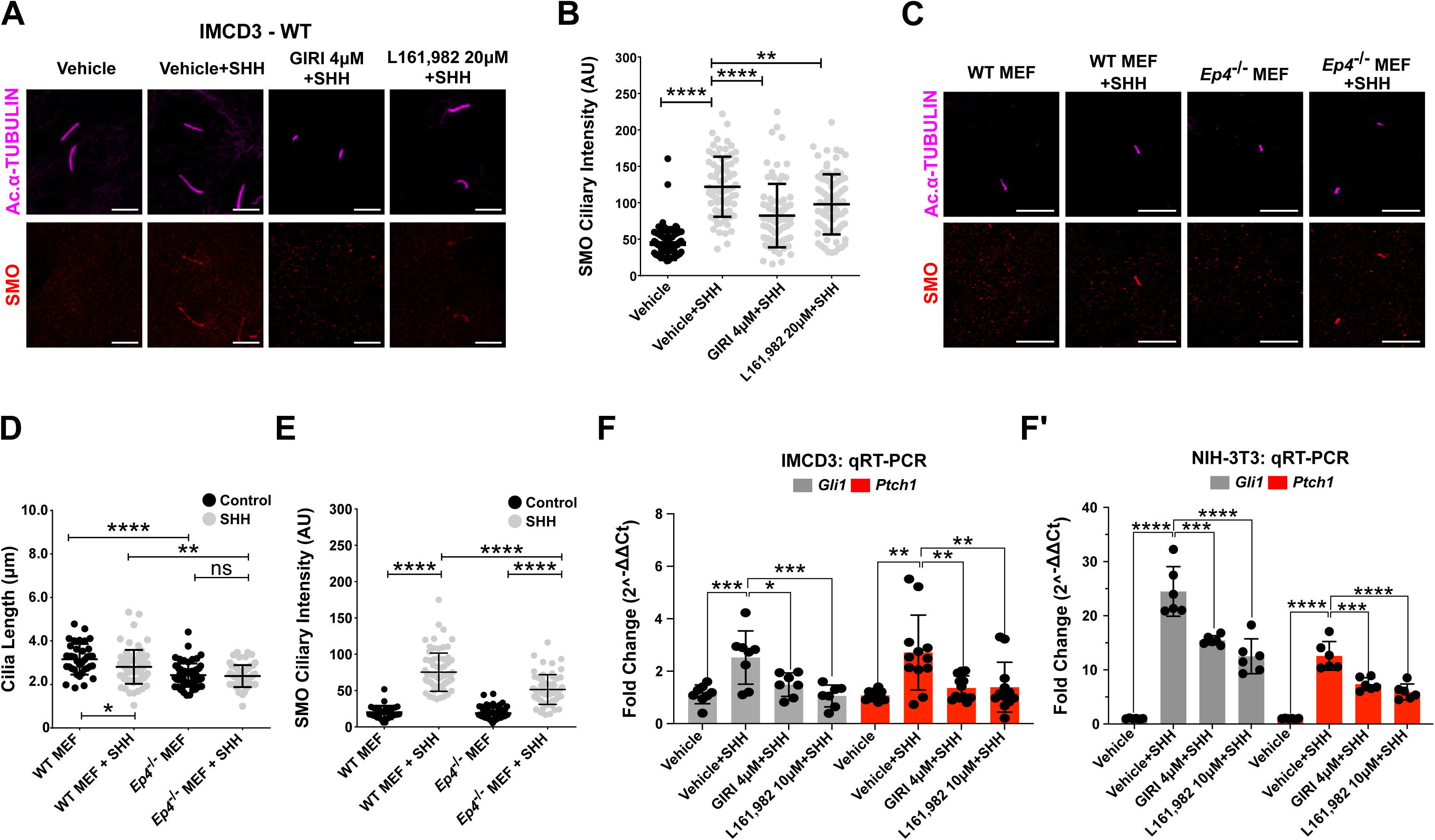
SHH signaling is attenuated by inhibition of cPLA_2_ or EP_4_. **(A-B).** SMO ciliary enrichment is reduced by EP_4_ inhibition. SMO (**A,** red) enrichment in primary cilia was examined in IMCD3 cells. Primary cilia are marked with acetylated α-tubulin (magenta). Cells were pretreated with vehicle (DMSO), GIRI (4 µM), or L161,982 (20 µM) for 18 hours. Scale bar = 5 μm. **(B).** Average SMO ciliary signal intensity was quantified for each condition. **(C-E).** EP_4_ loss shortens primary cilia and attenuates SMO ciliary enrichment. Ciliary length and SMO (red) ciliary enrichment were examined in wild type and *Ep4^-/-^* MEFs 18 hours post treatment with control or SHH conditioned media. Cilia are marked by acetylated α-tubulin (magenta, **C**). Scale bar =10 μm. A representative experiment is shown. Quantification of average primary cilium length **(D)** and ciliary SMO signal intensity **(E)** were determined as above by measuring cilia of ≥150 cells/condition across three independent experiments. **(F-F’).** qRT-PCR analyses of *Gli1* and *Ptch1* expression in IMCD3 and NIH-3T3 cells were performed. Cells were pretreated with vehicle (DMSO), GIRI (4 µM), or L161,982 (10 µM) for 2 hours and then cultured in control or SHH conditioned media plus inhibitor for 18 hours. Fold-change in expression was determined using the 2^−ΔΔCt^ method. Average fold change was calculated across two independent experiments with three biological replicates per experiment. All data are pooled. For all experiments, statistical significance was determined using a one-way ANOVA. A p value of less than 0.05 was considered statistically significant. Significance is denoted as follows: *<0.05, **<0.01, ***<0.001, ****<0.0001, and ns, p > 0.05. Error bars indicate SD.

### EP_4_ signaling stimulates ciliary cAMP recovery following SHH pathway activation

We hypothesized that the ciliary length reduction and SHH signal attenuation observed following EP_4_ inhibition resulted from altered ciliary cAMP regulation (Figure 4A). To address this hypothesis, we first tested whether reduced ciliary cAMP would lead to shortening of IMCD3 primary cilia by knocking down expression of ciliary adenylyl cyclases AC3, AC5, or AC6 (Supplementary Figure 2A-B). Consistent with these ACs localizing to primary cilia, their knockdown did not prevent forskolin-stimulated accumulation of whole cell cAMP by cytoplasmic ACs (Supplementary Figure 1C). To specifically measure IMCD3 cell ciliary cAMP following AC knockdown, we used a primary cilium-localized fluorescent sensor system to track ciliary cAMP changes in real time (Supplementary Figure 2D-F’). The sensor revealed that knockdown of each of the ciliary ACs attenuated ciliary cAMP increase by the AC stimulating drug forskolin (Figure 4B). Consistent with reduced ciliary cAMP leading to ciliary shortening, we observed a significant decrease in average IMCD3 cell primary cilium length following knockdown of *Adcy3*, *Adcy5*, or *Adcy6* (Figure 4C). Consistent with reduced primary cilium length compromising the SHH signal response, *Gli1* induction was attenuated following ciliary AC knockdown (Figure 4D), suggesting that sustained depletion of ciliary cAMP can compromise ligand-stimulated SHH pathway induction.

**Figure 4:**
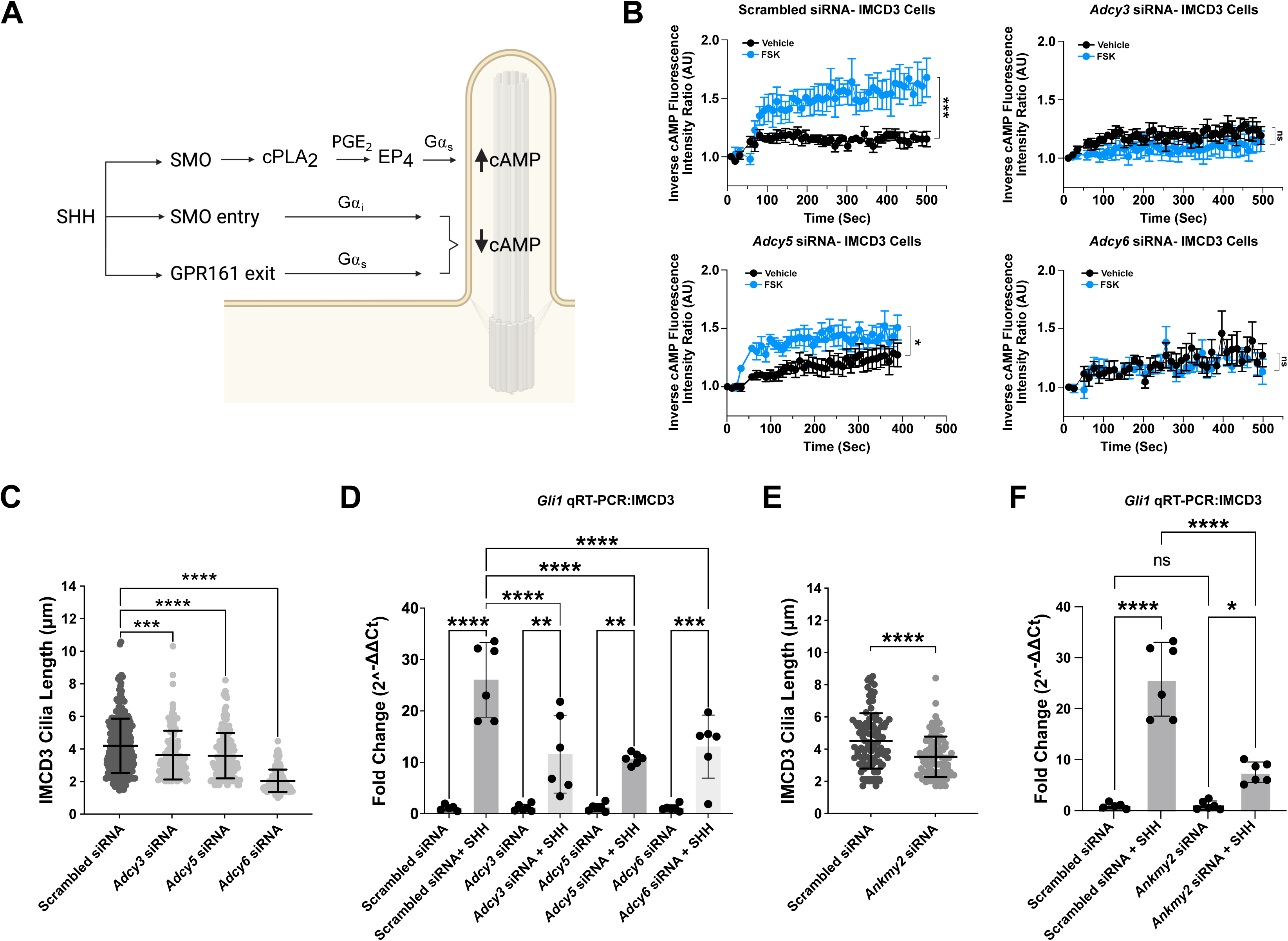
Ciliary adenylyl cyclases contribute to primary cilium length control and SHH stimulated transcriptional activation. **(A).** A model for SHH and EP_4_ effects on primary cilium cAMP control. **(B-D).** Knockdown of ciliary AC3, AC5, or AC6 disrupts ciliary cAMP control, shortens primary cilia, and compromises *Gli1* induction. **(B).** IMCD3 cells were treated with scrambled, *Adcy3*, *Adcy5,* or *Adcy6* siRNA for 48 hours and ciliary cAMP was measured using the cADDis cAMP sensor. cADDis activity was recorded as cells were exposed to vehicle (ethanol, black) or forskolin (FSK,100 μM, blue). Fluorescence ciliary intensity was recorded over 8 minutes in live cell imaging mode. An average of ∼5 cilia were recorded for each condition. The experiment was performed three times. A representative experiment is shown. Significance was determined by calculating the area under the curve followed by Student’s t-test analysis. **(C)**. Ciliary AC knockdown shortens primary cilia. Knockdown experiments were repeated at least twice with 75-100 primary cilia measured per experiment per condition. All data were pooled. Significance was determined by one-way ANOVA. **(D)**. *Gli1* fold change was determined in control and SHH-stimulated IMCD3 cells following *Adcy3*, *Adcy5*, or *Adcy6* knockdown. Fold change was assessed by qRT-PCR ∼48 hours after treatment with AC-specific siRNA or scrambled control following 2 hours of serum starvation and overnight treatment with SHH conditioned media. Expression was determined using the 2^−ΔΔCt^ method. Significance was determined by a one-way ANOVA. **(E-F)**. Knockdown of *Ankmy2* shortens primary cilia and attenuates ligand-stimulated *Gli1* induction. **(E).** Knockdown experiments were repeated at least twice with 75-100 primary cilia measured per experiment per condition. All data were pooled. Significance was determined by Student’s t-test (F). *Gli1* fold change was determined in control or SHH-stimulated IMCD3 cells following *Ankmy2* knockdown, as described above. The experiment was performed twice in triplicate and all data pooled. Significance was determined by a one-way ANOVA. For all experiments, a p value of less than 0.05 was considered statistically significant. Significance is denoted as follows: *<0.05, **<0.01, ***<0.001, ****<0.0001, and ns, p > 0.05. Error bars indicate SD.

Despite the established link between ciliary length destabilization and reduced SHH pathway activation (Caspary et al., 2007; He et al., 2014), our observation that decreased ciliary cAMP correlated with reduced *Gli1* induction was curious. This is because reduced ciliary AC localization following loss of the ciliary AC chaperone protein ANKMY2 has been shown to induce GLI gain-of-function phenotypes *in vivo* (Somatilaka et al., 2020). To query this discrepancy, we knocked down *Ankmy2* in IMCD3 cells to probe effects of combined ciliary AC depletion on primary cilium cAMP concentration and length control. Consistent with published results, *Ankmy2* knockdown reduced ciliary enrichment of endogenous AC3 and AC5/6 (Somatilaka et al., 2020) and blocked the ability of forskolin to trigger rapid ciliary cAMP accumulation (Supplementary Figure 3A-D). To determine how ciliary AC depletion impacted primary cilium length and SHH pathway induction, we measured primary cilia and performed *Gli1* qPCR analyses in control and *Ankmy2* knockdown cells treated with control or SHH conditioned media. Notably, *Ankmy2* knockdown triggered a 23% reduction in IMCD3 cell primary cilium length and attenuated SHH-stimulated *Gli1* induction (Figure 4E-F). These results suggest the inability of cells to raise ciliary cAMP in SHH-stimulated cells compromises the ligand-induced transcriptional response.

To further evaluate this hypothesis, we next probed effects of ciliary AC depletion on the SHH response in highly SHH-responsive NIH-3T3 cells. Due to baseline primary cilia lengths in NIH 3T3 cells being shorter than those of IMCD3 cell cilia, length reductions occurring in response to ciliary AC depletion were difficult to appreciate. Nevertheless, we observed that knockdown of *Adcy5* or *Ankmy2* led to modest, but statistically significant, reduction in primary cilium lengths in NIH-3T3 cells (Supplementary Figure 3E-F). *Adcy3* knockdown did not alter NIH-3T3 ciliary length, suggesting AC5 may be the primary modulator of cAMP-mediated primary cilium length regulation in this cell type (Supplementary Figure 3F). Accordingly, and consistent with published results (Somatilaka et al., 2020), we observed that knockdown of *Ankmy2* significantly reduced SHH-stimulated *Gli1* induction in NIH-3T3 cells (Supplementary Figure 3G). Consistent with *Ankmy2* knockdown leading to a pan-AC ciliary depletion, individual knockdown of either *Adcy3* or *Adcy5* attenuated the SHH transcriptional response to a lesser extent than *Ankmy2* knockdown (Supplementary Figure 3G). We did not detect endogenously expressed *Adcy6* transcript in NIH-3T3 cells, so did not test knockdown of *Adcy6* in this cell type. These results, combined with the results presented above, lead us to conclude that primary cilium AC depletion is detrimental to the ligand-induced SHH signal response.

Having established that lowering ciliary cAMP by AC depletion appreciably and significantly shortened primary cilia and attenuated SHH signaling in IMCD3 and NIH-3T3 cells, we next sought to determine whether cAMP dynamics downstream of SMO activation were impacted by EP_4_. Thus, IMCD3 cells were stimulated with the AC activator forskolin, and ciliary cAMP was monitored in control and SMO stimulated cells using the fluorescent cAMP system introduced above. As anticipated, forskolin treatment stimulated a rapid increase in ciliary cAMP fluorescence intensity ratios (Figure 5A, red and Supplementary Figure 2F-F’, blue). Consistent with the ability of activated SMO to couple with Gα_i_ and induce exit of Gα_s_-coupled GPR161 from primary cilia, co-stimulation of IMCD3 cells with forskolin and the direct SMO agonist SAG reduced the level to which forskolin could raise ciliary cAMP. Despite this, ciliary cAMP in cells treated with forskolin + vehicle and forskolin + SAG equilibrated similarly after an initial lag (Figure 5A, red vs blue). We hypothesized cAMP equilibration was the result of EP_4_ activation downstream of SMO-stimulated PGE_2_ release. Accordingly, ciliary cAMP equilibration in SAG stimulated cells was blocked by direct EP_4_ inhibition with L161,982 (Figure 5A, green vs. blue). We conclude EP_4_ activation is required to equilibrate primary cilium cAMP levels in cells with an active SHH signaling pathway.

**Figure 5:**
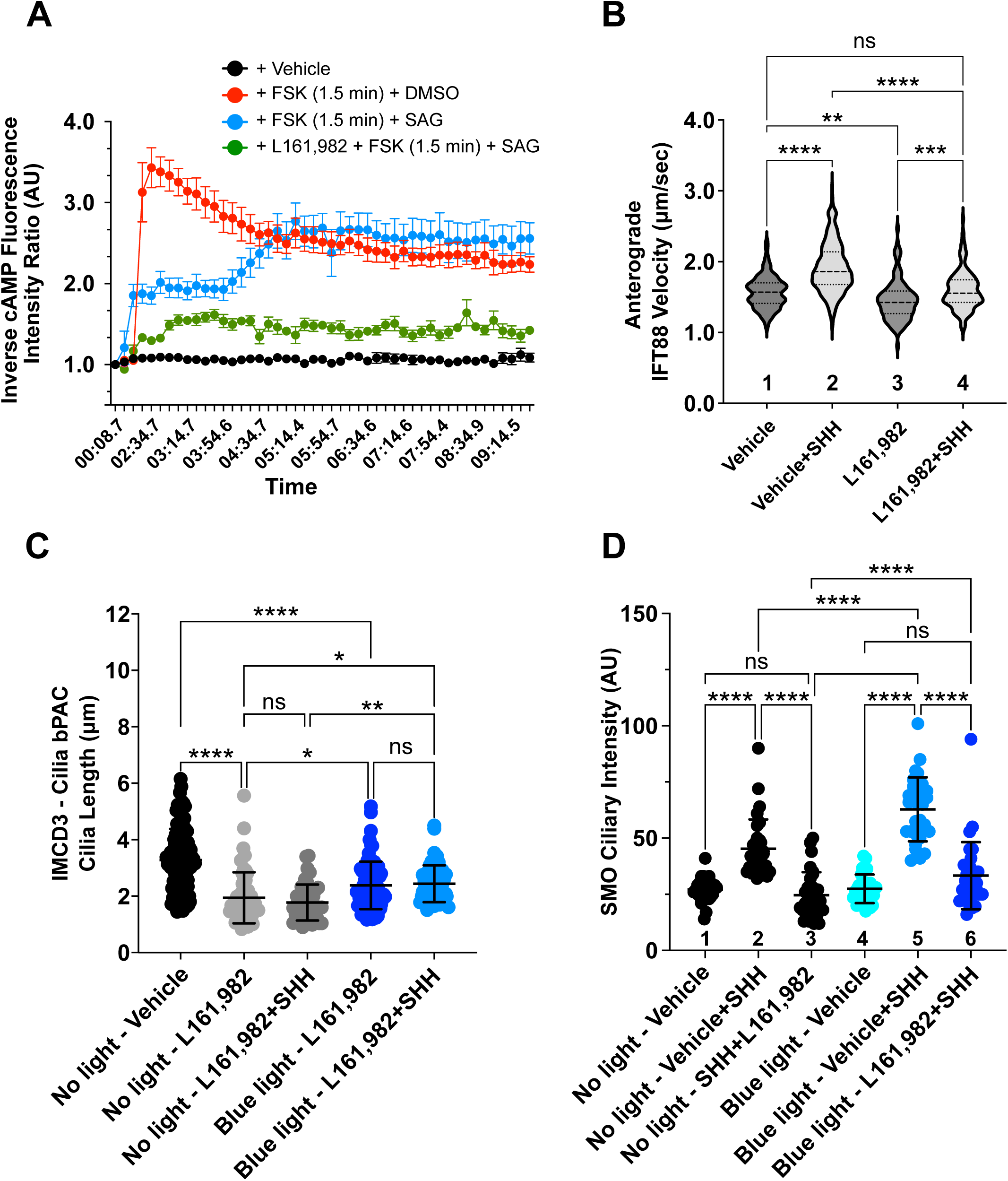
EP_4_ stabilizes ciliary cAMP in SHH-stimulated cells to maintain anterograde IFT and SMO ciliary accumulation. **(A).** EP_4_ signaling equilibrates SMO-induced reduction of primary cilium cAMP. cADDis-expressing IMCD3 cells were pretreated overnight with vehicle or L161,982 (10 µM). The following morning, cADDis activity was monitored by live imaging to track the ciliary cAMP response as cells were treated with forskolin (FSK, 100 μM) for 1.5 minutes prior to addition of the SMO agonist SAG (1 μM) or vehicle control. Fluorescence ciliary intensity was recorded over 10 minutes in live cell imaging mode. An average of ∼6 cilia were recorded for each condition and the experiment was performed twice. A representative experiment is shown. **(B)**. Anterograde IFT velocity was calculated in IMCD3 cells by tracking IFT88-GFP movement in the presence and absence of SHH, L161,982 (10 µM), or vehicle control. IFT velocity was calculated for 30 cilia per condition across 4 experiments. Velocity is shown as a violin plot with SD indicated. **(C-D).** Average ciliary length and SMO ciliary intensity were quantified in IMCD3-bPAC cells exposed to control or SHH conditioned media in the absence or presence of 10 µM L161,982 in control or blue light-exposed cells. Significance was determined by one-way ANOVA. For all experiments, a p value of less than 0.05 was considered statistically significant. Significance is denoted as follows: *<0.05, **<0.01, ***<0.001, ****<0.0001, and ns, p > 0.05. Data are represented as mean ± SD. For all ciliary analyses, 50-100 cells per condition were analyzed over at least 3 independent experiments.

Previous reports indicate that activation of EP_4_ raises cAMP to promote anterograde IFT (Besschetnova et al., 2010; Jin et al., 2014). To determine if this activity was compromised by EP_4_ inhibition in SHH-stimulated cells, we calculated anterograde velocity of the ciliary intraflagellar transport protein IFT88 in control and L161,982-treated cells. We observed increased anterograde IFT in vehicle-treated, SHH-stimulated cells (Figure 5B, lane 1 vs. 2). Strikingly, anterograde IFT88 velocity was significantly reduced following treatment with the EP_4_ inhibitor L161,982 in both control and SHH-stimulated IMCD3 cells (Figure 5B). Thus, EP_4_ activity is required to maintain anterograde IFT in control cells and enhance its velocity in SHH stimulated cells.

If cAMP equilibration that stabilizes primary cilium length in SHH-stimulated cells occurs through SHH-to-EP_4_ crosstalk, we reasoned that providing supplemental ciliary cAMP might restore ciliary length to rescue SHH pathway induction in L161,982-treated cells. To affect ciliary cAMP in a controlled manner, we adapted a published optogenetic system (Truong et al., 2021) for use in IMCD3 cells. In this system, bacterial photoactivatable adenylyl cyclase (bPAC) is expressed in the cell cytoplasm or targeted to primary cilia through fusion with the primary cilium-localized protein ARL13B, and then activated to specifically raise cytoplasmic or ciliary cAMP by blue light stimulation (Supplementary Figure 4A). IMCD3 cells were engineered to stably express cytoplasmic or ciliary bPAC-GFP, and then exposed to blue light to quantify primary cilium length changes (Supplementary Figure 4B-C”). IMCD3 cells expressing cytoplasmic or ciliary bPAC proteins showed similar average primary cilium lengths in the absence of blue light and cilia of cyto-bPAC-expressing cells did not change length upon blue light exposure, despite its ability to increase cytoplasmic cAMP levels (Supplementary Figure 4C-C’). Conversely, cilia-bPAC-expressing cells increased their average primary cilium length by ∼30% upon blue light exposure (Supplementary Figure 4C, C”). To support that elongation resulted from increased primary cilium cAMP, we measured primary cilium lengths in IMCD3 cells treated with forskolin, which stimulates both ciliary and non-ciliary AC. We observed an increase in average primary cilium length in forskolin-treated cells that was similar to that observed following cilia-bPAC activation (Supplementary Figure 4D-D’). Thus, small molecule or optogenetic increase in ciliary cAMP can increase primary cilium length.

Having validated feasibility of using ciliary bPAC to modulate primary cilium cAMP in IMCD3 cells, we next determined whether shortened cilia of EP_4_-inhibited cells could be reversed by increasing ciliary cAMP. Consistent with the hypothesis that reduced primary cilia lengths in SHH-stimulated L161,982-treated cells resulted from failed cAMP equilibration, blue light stimulation of cilia-bPAC partially rescued average primary cilium length in EP_4_-inhibited, SHH stimulated cells. Blue light stimulation increased average ciliary lengths of L161-982-treated, ciliary-bPAC-expressing cells by ∼25% in the absence and presence of SHH (Figure 5C). To test whether ciliary length increase correlated with rescue of SHH pathway activity, we quantified SMO ciliary enrichment in control and blue light-stimulated, cilia-bPAC-expressing cells after L161,982 treatment. Consistent with the ability of blue light-induced cAMP to partially rescue ciliary length following EP_4_ inhibition, SHH-stimulated SMO ciliary enrichment was also partially restored in blue light-stimulated, L161,982-treated cilia-bPAC cells (Figure 5D, column 3 vs. 6). Intriguingly, SHH-stimulated enrichment of SMO in vehicle-treated cells was enhanced by blue light stimulation, supporting that augmenting cAMP production in cells with an active SHH pathway may enhance anterograde IFT to facilitate increased SMO ciliary enrichment (Figure 5D, column 2 vs. 5). Thus, EP_4_-regulated ciliary cAMP control impacts both primary cilium length and SMO ciliary accumulation.

### EP_4_ is required for ciliary length control and SHH signaling during neural tube development

Having established a cell biological connection between SHH and EP_4_ that stabilizes primary cilium length, we next sought to determine whether EP_4_ loss compromised ciliary length control to impact SHH signaling *in vivo*. Thus, we analyzed developing neural tubes of *Ep4^-/-^* mice to assess primary cilium length and evaluate SHH-regulated ventral neural tube cell fate specification. *Ep4^-/-^* mice are present at normal Mendelian ratios *in utero* but die as neonates due to heart failure resulting from highly penetrant patent ductus arteriosus (Segi et al., 1998). Thus, we collected E9.5/29 somite stage *Ep ^+/-^* and *Ep ^-/-^* embryos to perform scanning electron microscopy for visualization of apical primary cilia of cells lining the neural tube lumen (Figure 6A). Measurement of apical primary cilia revealed statistically significant length reductions in *Ep4^-/-^* embryos compared to cilia of heterozygous littermate controls (Figure 6B). Whereas primary cilia in control neural tubes averaged ∼0.9 µm, cilia in knockout embryos showed an approximate 30% length reduction to an average length of ∼0.6 µm. Thus, EP_4_ contributes to primary cilium length control in the developing neural tube.

**Figure 6:**
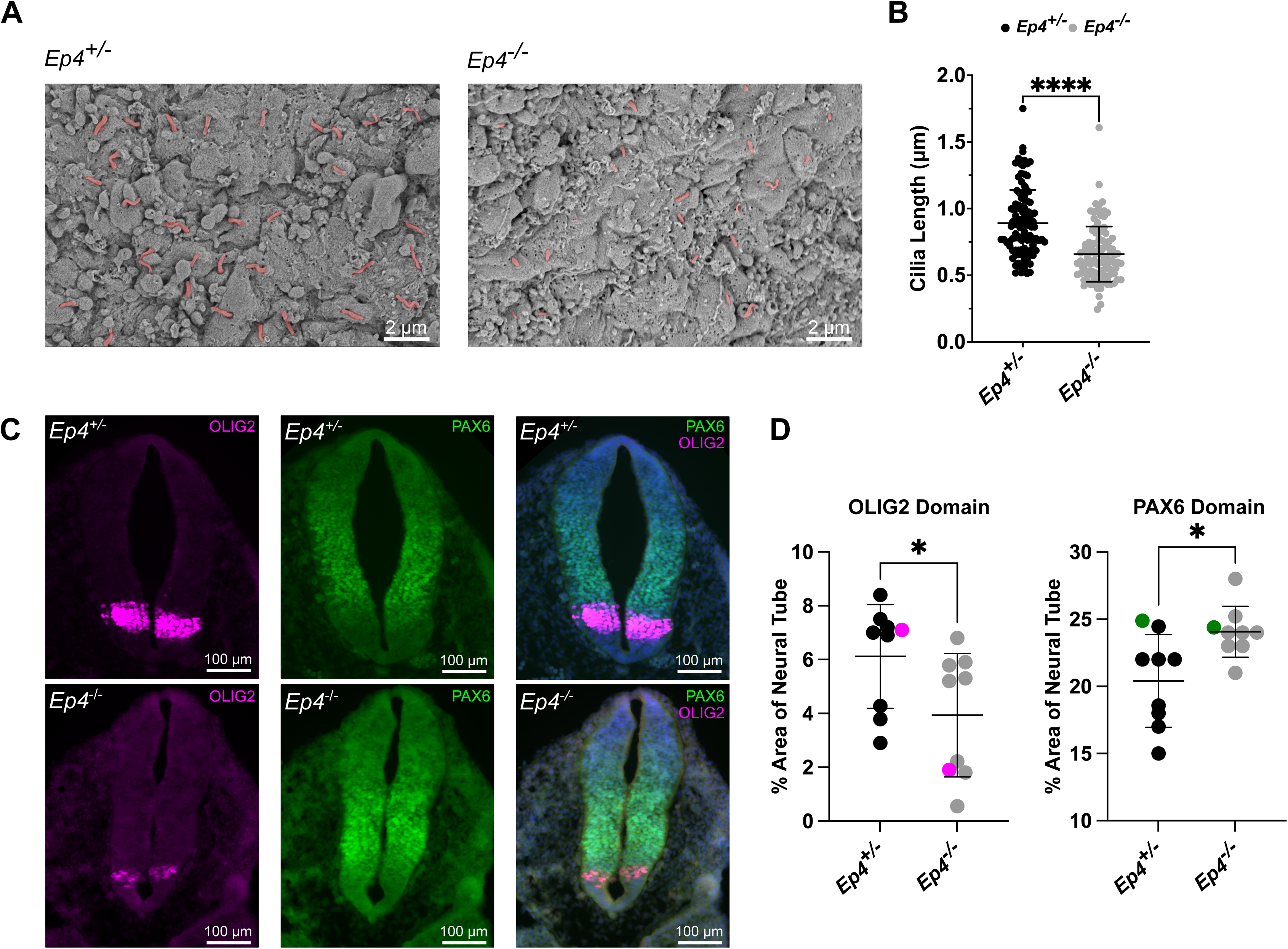
EP_4_ loss *in vivo* reduces primary cilium length and alters SHH-dependent neural tube patterning. **(A)**. Scanning electron microscopy images of cardiac level neural tube sections from E9.5/27 somite stage *Ep4^+/-^* and *Ep4^-/-^* embryos show apical plasma membranes of cells lining the neural tube lumen. Scale bar = 2 μm. Cilia are shaded in pink **(B)**. Average primary cilium length was calculated by measuring cilia of ∼30 cells/embryo lining the neural tube lumen. Primary cilia of *Ep ^-/-^* embryos show a statistically significant length reduction. Statistical significance was calculated using one-way ANOVA. Error bars indicate SD. Significance is denoted as follows: ****<0.0001, and ns, p > 0.05. **(C-D).** *Ep ^-/^*^-^ embryos show altered ventral progenitor domains. Cardiac level sections of E9.5/29 somite stage neural tubes from the indicated genotypes were immuno-stained for neural progenitor domain markers *Olig2* (magenta) and *Pax6* (green). *Ep ^-/-^* neural tubes show disrupted ventral progenitor specification as indicated by reduced *Olig2*-labeled cellular populations. For all experiments, four embryos per genotype were analyzed. Scale bar = 100 μm. Progenitor domain area is shown in **(D)**. Statistical significance was calculated using a Student’s t-test. Error bars indicate SD. Significance is denoted as follows: *<0.05. Sections shown in **(C)** are indicated as colored dots in each graph.

During neural tube development, SHH signals in a graded manner from the notochord and neural tube floor plate to instruct transcriptional programs that establish ventral neural tube cell fates (Ericson et al., 1997; Patten and Placzek, 2000; Ribes and Briscoe, 2009). To test whether EP_4_ loss led to alteration of SHH-dependent neural tube progenitor domain specification, we examined expression of the ventral SHH transcriptional target *Olig2,* which is activated by high SHH, and the intermediate fate marker *Pax6* (Ericson et al., 1997; Ribes and Briscoe, 2009).

Because specification of ventral progenitor domains is highly sensitive to SHH signal alteration, monitoring their induction allows for reliable reporting of SHH signaling activity (Ribes and Briscoe, 2009). Comparison of E9.5/27 somite stage knockout and heterozygous littermate control neural tubes revealed that EP_4_ loss led to reduction in the *Olig2*-postive motor neuron progenitor cell population, indicative of reduced SHH signaling response in *Ep4^-/-^* mice.

Coincident with reduction in the *Olig2*-expressing progenitor domain, we observed expansion of the *Pax6*-expressing progenitor population (Figure 6C-D). Taken together with the above, these results support that EP_4_ loss shortens primary cilia and compromises SHH-directed cell fate specification *in vivo*.

## DISCUSSION

Prostaglandin signaling is important for motile cilia development and beat frequency and contributes to primary cilium elongation (Gayner and McCaffrey, 1998; Haxel et al., 2001; Jin et al., 2014; Marra et al., 2019). The study presented here reveals the SHH pathway, which requires functional primary cilia to initiate downstream effector responses, links with the PGE_2_ pathway to ensure ciliary length homeostasis. We delineate a SHH-to-PGE_2_/EP_4_ signal crosstalk circuit and demonstrate that SHH-stimulated PGE_2_ production leads to activation of Gα_s_-coupled EP_4_. EP_4_ signaling equilibrates ciliary cAMP after an initial dip following SHH/SAG stimulated ciliary exit of Gα_s_-coupled GPR161 and activation of Gα_i_-coupled SMO. We propose that cAMP equilibration maintains anterograde IFT that is necessary for SMO ciliary enrichment and primary cilium length stability (Figure 7A-B). Genetic or pharmacological interruption of the SHH-to-EP_4_ signal crosstalk cascade short-circuits ciliary cAMP equilibration in SHH-stimulated cells, slows anterograde IFT, shortens primary cilia, and attenuates SHH signal output (Figure 7C). *In vivo* loss of EP_4_ leads to short neuroepithelial primary cilia and neural tube defects indicative of compromised SHH signaling. These results support an essential link between SHH and EP_4_ ciliary signaling pathways during embryonic development.

**Figure 7:**
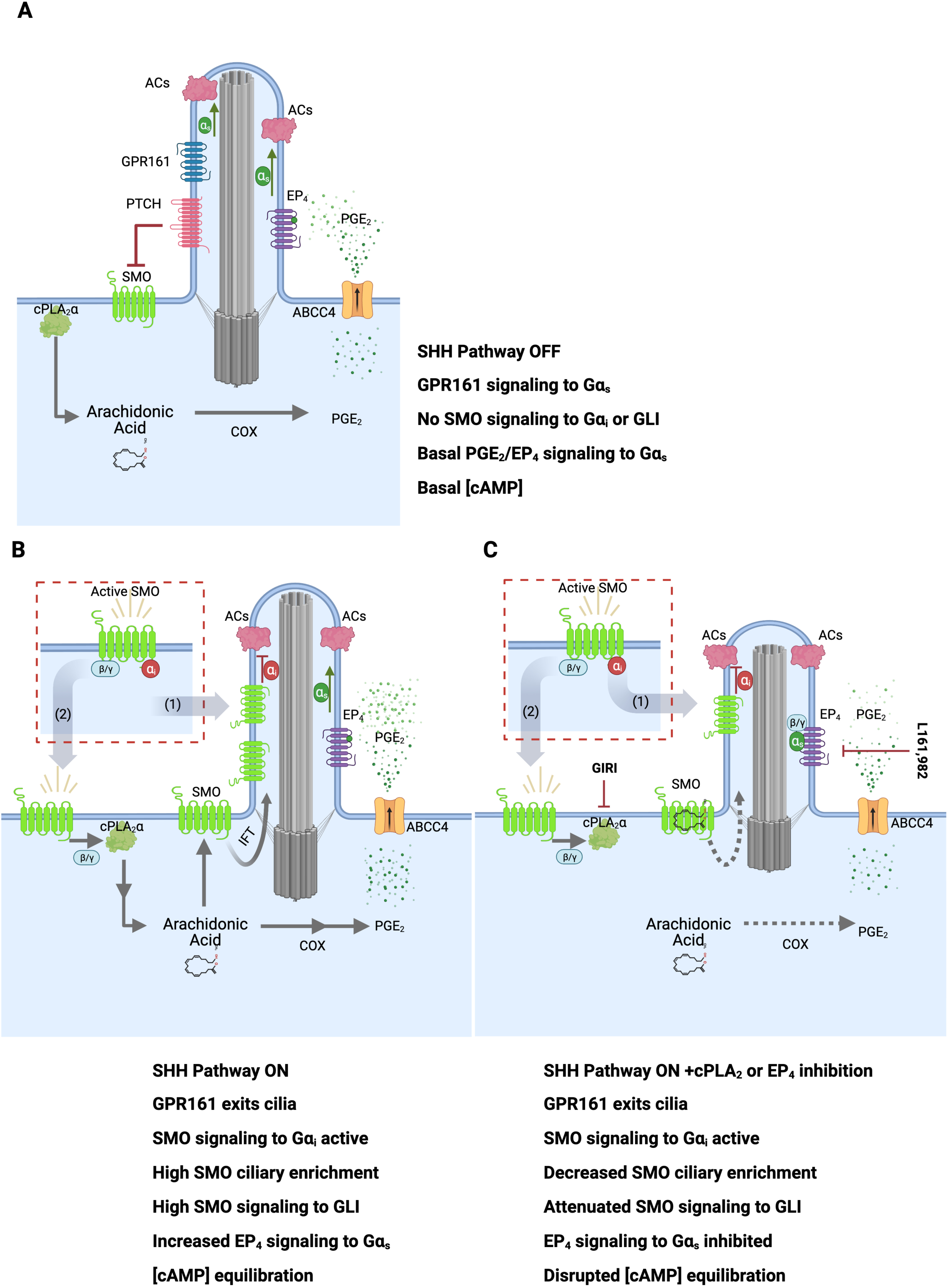
A model for SHH-to-EP4 signal crosstalk. **(A).** In the absence of SHH, the SHH receptor PTCH prevents ciliary accumulation and signaling of the GPCR SMO. In this basal condition, GPR161 signaling and basal PGE_2_ secretion and activation of EP_4_-Gα_s_ maintain sufficient ciliary cAMP for anterograde IFT and length homeostasis. **(B).** SHH binding to PTCH leads to SMO de-repression and GPR161 ciliary exit. SMO activates Gα_i_βγ to increase production of arachidonic acid by βγ stimulation of cPLA_2_α. Arachidonic acid is metabolized to PGE_2_, which is secreted to activate ciliary EP_4_ for Gα_s_ activation. This ensures ciliary cAMP levels remain sufficiently high in SHH-stimulated cells to prevent anterograde IFT slowing that leads ciliary shortening. **(C).** Inhibition of cPLA_2_ or EP_4_ reduces SHH-to-EP_4_ signal crosstalk, reduces primary cilium length, and blunts SMO ciliary accumulation and high-level signaling.

Although the requirement for functional primary cilia in regulation of SHH signaling is well established, the complex relationship between SHH and cAMP modulation in cilia has yet to be clearly defined. To initiate a transcriptional response, SHH must lower ciliary cAMP to halt PKA instructed GLI processing into its repressor form (Mukhopadhyay et al., 2013; Niewiadomski et al., 2014). Consistent with this biology, genetic loss of the AC chaperone *Ankmy2* depletes primary cilia of AC activity, attenuates PKA-directed GLI repressor conversion, and allows for unregulated accumulation of full length GLI proteins that induce gain-of-function phenotypes *in vivo*. Although ciliary length reduction was not noted in *Ankmy2* knockout mice, mechanistic *in vitro* analyses revealed that *Ankmy2* knockout NIH-3T3 cells were, nonetheless, compromised in their ability to induce SHH-stimulated transcriptional responses (Somatilaka et al., 2020). This suggests the ability to precisely tune ciliary cAMP levels up or down is essential for optimal ligand-stimulated SHH pathway activity. Our data suggest SHH-promoted EP_4_ signaling is a key mechanism by which up-tuning occurs, and that this crosstalk is necessary to prevent slowing of anterograde IFT and primary cilium shortening that can occur in response to sustained ciliary cAMP depletion. However, the requirement for SHH to activate Gα_s_-coupled EP_4_ to stabilize primary cilium length prompts the question of how GLI escapes PKA-mediated repressor conversion upon ciliary cAMP re-elevation. We hypothesize GLI stabilization is maintained in SHH-stimulated cells through direct PKA inhibition by active SMO. Recent reports demonstrate that SHH-stimulated phosphorylation of the intracellular carboxyl-terminal tail of SMO exposes a PKA inhibitory domain (PKI) that prevents GLI phosphorylation (Arveseth et al., 2021; Happ et al., 2022). As such, we propose SHH coordinates a sophisticated ciliary signal response through a network of interconnected GPCR activities.

Notably, GPCR signal crosstalk downstream of Hedgehog (Hh) family ligands that equilibrates cAMP may be a conserved mechanism by which pathway activity is controlled. In *Drosophila*, Hh-activated Smo stimulates Gα_i_ to lower cAMP and halt PKA-regulated truncation of the GLI ortholog Cubitus interruptus (Ci) (Ogden et al., 2008). However, a *Drosophila*-specific positive regulatory role for PKA-mediated phosphorylation of Smo is required for high-level pathway induction, and could be disrupted by Smo activation of Gα_i_ (Jia et al., 2004). A potential mechanism by which Smo solves this problem was revealed by the discovery that, in some cell types, Smo stimulation leads to Gα_s_ activation to enhance pathway responsiveness to low threshold Hh stimulation (Praktiknjo et al., 2018). It is not yet clear whether Gα_s_ activation downstream of Hh occurs through Smo signal bias or by Smo crosstalk with Gα_s_-coupled GPCR signaling partners. Nevertheless, the observation that Smo can lead to Gα_s_ activation in flies supports an evolutionarily conserved ability of Hedgehog family members to affect exquisite control of cAMP modulators to ensure appropriate downstream signaling activity.

**Table 1:**
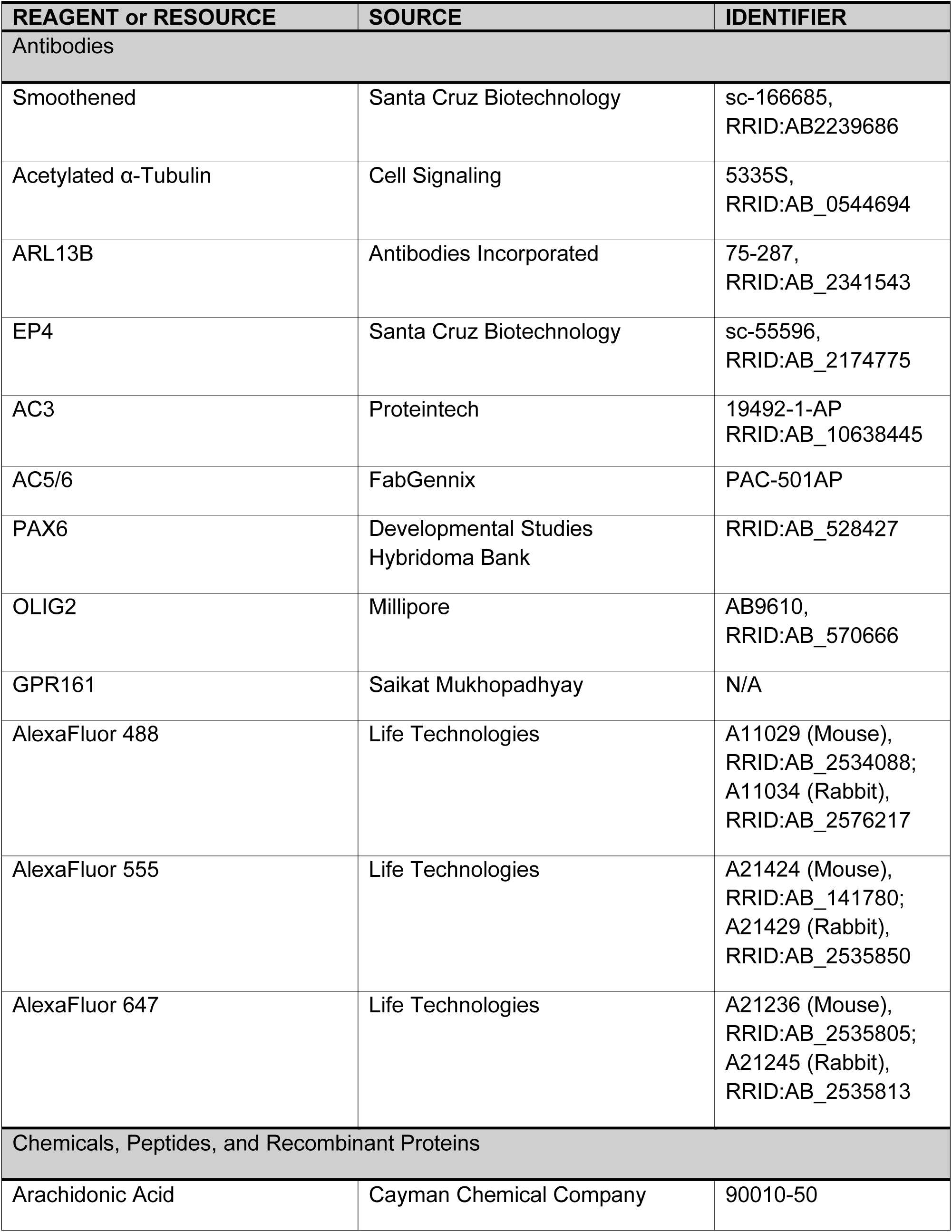
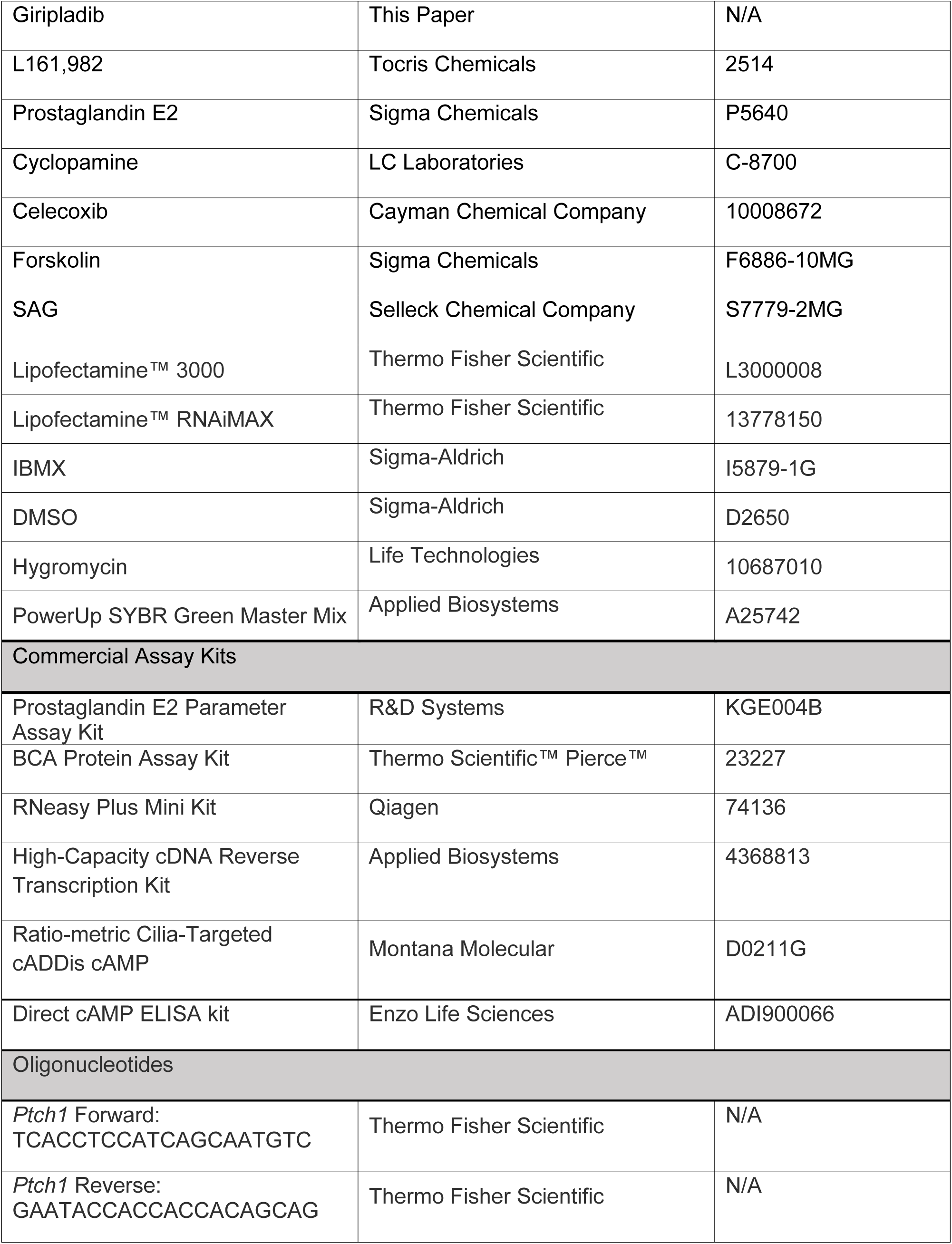
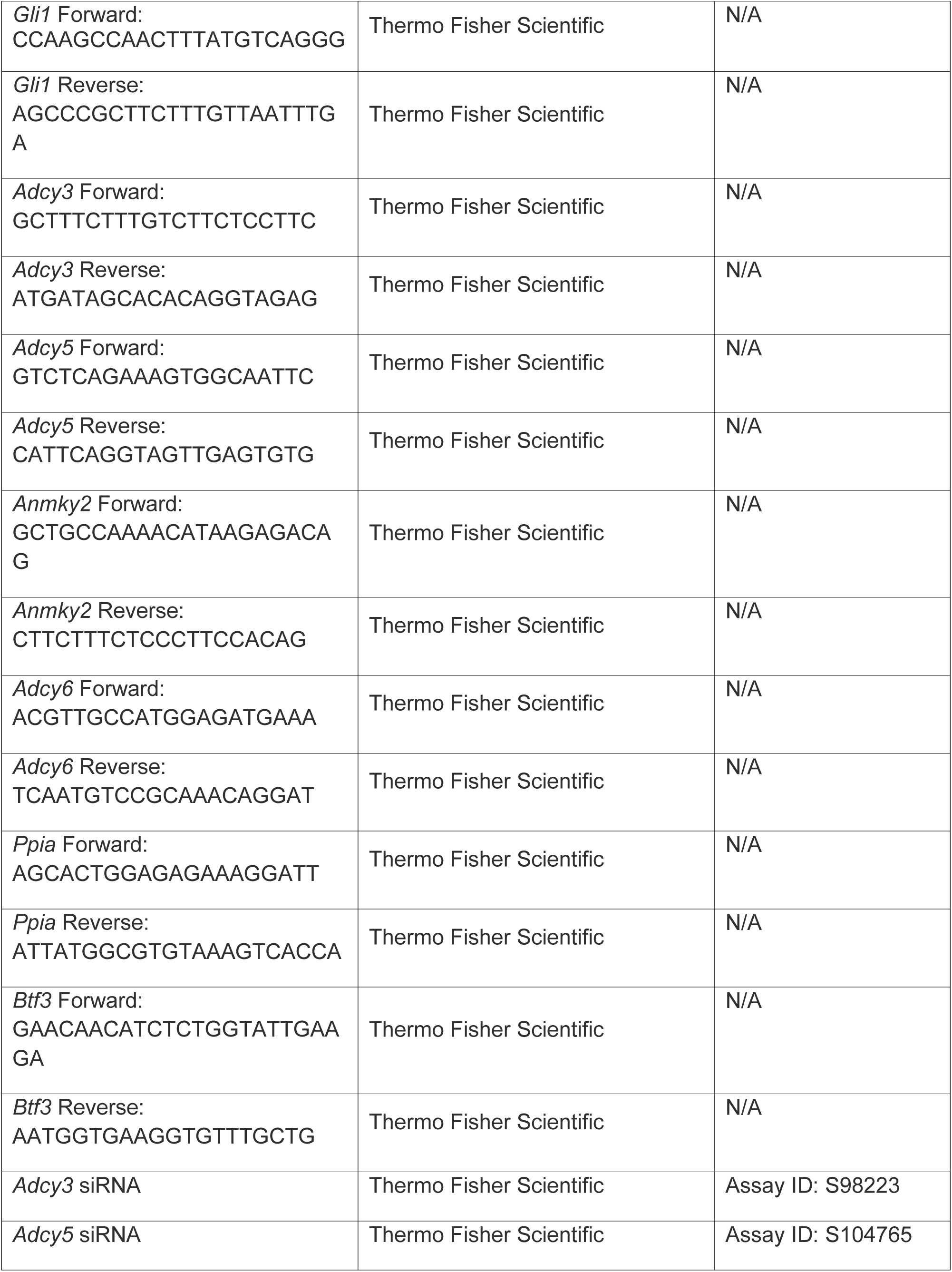
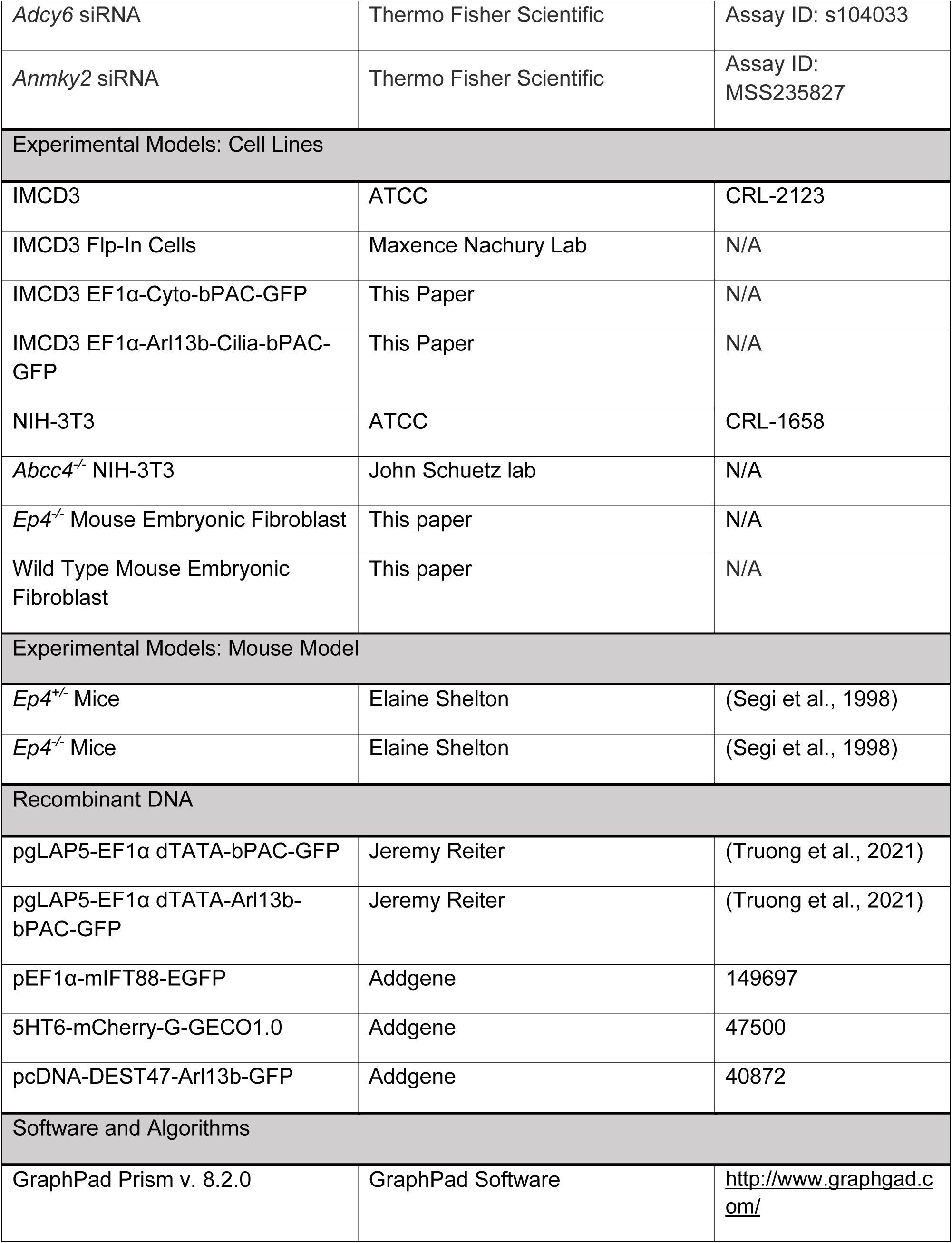
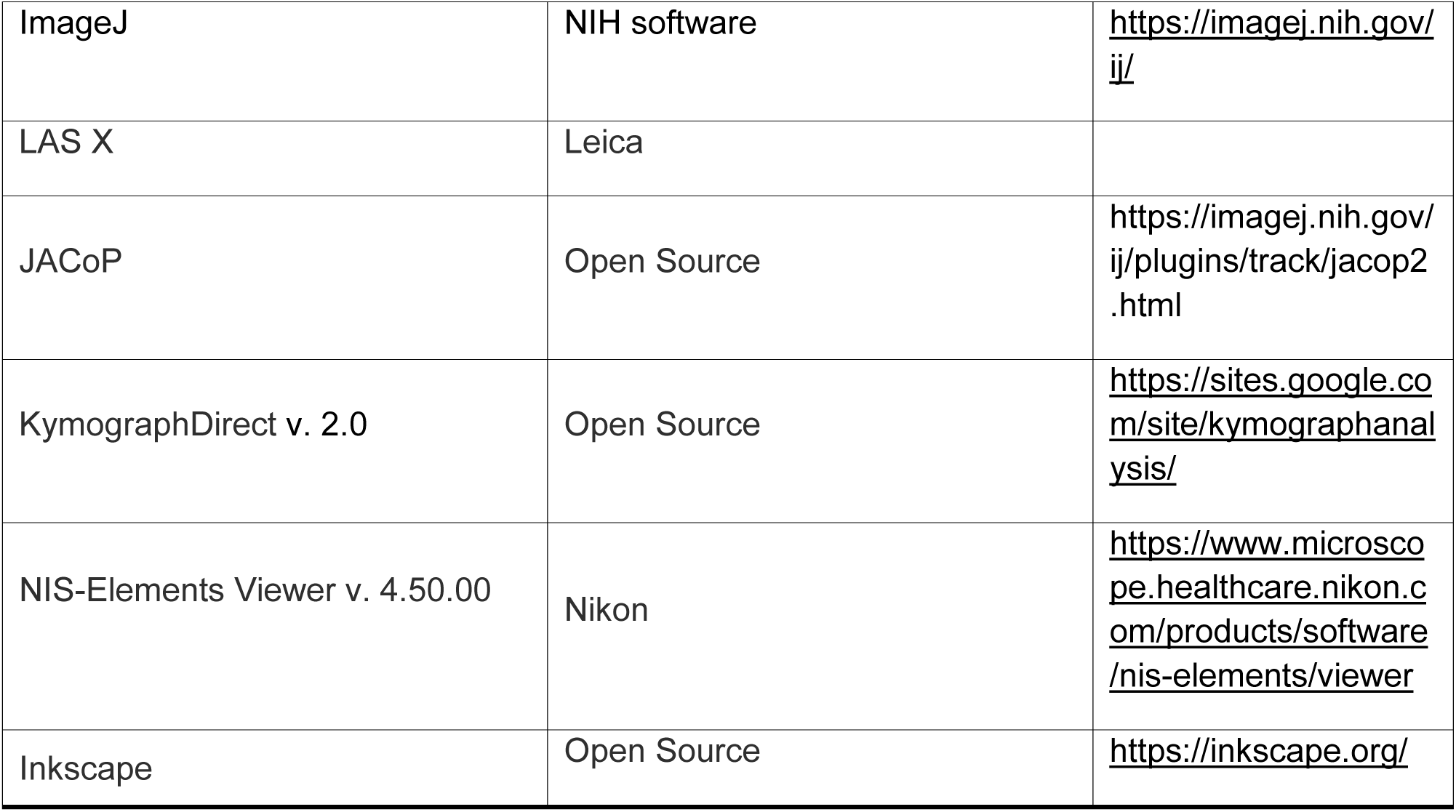
Reagents and Resources

## RESOURCE AVAILABILITY

Requests for resources and reagents should be directed to Stacey K. Ogden. Cell lines and novel reagents will be available upon request pursuant to SJCRH material transfer agreement.

## METHODS

### Mammalian Cell culture

Wild type and Flp-In IMCD3 cells were cultured in Dulbecco’s Modified Eagle Medium (DMEM): F12 supplemented with 10% fetal bovine serum (FBS) and 1% Penicillin Streptomycin solution (Gibco). Wild type MEFs, NIH-3T3, *Ep4*^-/-^ MEF, and *Abcc4^-/^*^-^-NIH-3T3, were supplemented with 10% fetal calf serum. Cells were passaged routinely with 0.25% Trypsin/EDTA solution and maintained under standard incubation conditions (5% CO_2_, 37°C) in a humidified incubator. All cell lines were routinely tested for mycoplasma contamination. To promote ciliogenesis, cell lines were serum starved for 2 hours in serum-free media (DMEM or DMEM: F12, supplemented with 0.1 mM nonessential amino acids, 2mM L-Glutamine, 1mM sodium pyruvate, and 1% Pen-Strep). After starvation, media was replaced with low serum (0.5%) media and cells were incubated for 18 or 36 hours.

### Chemicals

L161,982 (Tocris), Cyclopamine (LC Laboratories) and Celecoxib (Coxib, Cayman Chemical Company) were reconstituted in DMSO and stored at -20°C. Arachidonic acid (Cayman Chemical Company) was stored in ethanol at -20°C. Prostaglandin E_2_ (Sigma Chemicals) and Forskolin (Sigma Chemicals) were reconstituted in ethanol and stored at 4°C. Giripladib (GIRI) was synthesized and stored according to published procedures (Duvernay et al., 2015; Thotala et al., 2013).

### Immunofluorescence and ciliary length measurement

For cultured cells ciliary analysis, cells were plated onto coverslips (Corning) and pretreated with either vehicle or drug in serum-free media. After 2 hours, media was changed to low serum media containing drug and SHH-N conditioned media (100 µL/mL), and then incubated for ∼18 hours. After incubation, cells were washed with PBS, and then fixed in 4% paraformaldehyde for 12 minutes. Cells were then washed with wash buffer (PBS with 0.1% Triton X-100) three times for 5 minutes and incubated in blocking buffer for 60 minutes at room temperature (PBS with 2% BSA, 0.1% Triton X-100, 1% goat serum). Primary antibody incubations were performed overnight at 4°C. The following antibodies and dilutions were used: anti-SMO (Santa Cruz, 1:500), anti-acetylated α-tubulin (Cell Signaling, 1:1000), anti-ARL13B (Antibodies Incorporated, 1:500), anti-EP_4_ (Santa Cruz, 1:100), anti-AC5/6 (FabGennix, 1:75), anti-AC3 (Proteintech, 1:200) diluted in blocking buffer. Secondary antibody incubations were performed with AlexaFluor 488, 555, or 647 conjugated secondary antibodies (Life Technologies, 1:1000) and DAPI for 60 minutes at room temperature. Following antibody incubation, samples were washed three times in wash buffer followed by a rinse in PBS, prior to mounting with ProLong Diamond (Life Technologies). Images were collected using Leica TCS SP8 STED 3x confocal microscope with a 63x oil-immersion objective and processed using LAS X (Leica). For all immunofluorescence experiments, multiple cells (≥100) were examined over at least two independent experiments. Representative images are shown.

For ciliary length measurement, a maximum projection of the acquired Z stack images was created. Each cilium was manually traced from the base to the tip using the line profile tool in the LAS X software. For all experiments, >100 cilia were measured for each condition across a minimum of two independent experiments. For quantification graphs, all data are included. For microscopy images, representative cilia are shown.

### In vivo analysis

Data and materials generated from animals were obtained in accordance with SJCRH/IACUC approved protocol 608-100616-10/19 and Vanderbilt protocol M1600016-01. All animal husbandry and procedures were performed in accordance with protocols approved by St. Jude Children’s Research Hospital and Vanderbilt University Institutional Animal Care and Use Committees.

*Ep4^+/-^* and *Ep4^-/-^* embryos in the C57BL/6 background were harvested and processed for immunohistochemistry at E9.5. Pregnant dams were harvested, uterine horns removed, and embryos were dissected in 1X PBS, then rinsed three times in 1X PBS. Embryos were fixed overnight at 4°C in 4% PFA. The following day, embryos were rinsed three times in 1X PBS and transferred to 30% sucrose to cryo-protect. The following day, embryos were frozen in O.C.T. Compound (Tissue-Tek) on dry ice. Embryos were sectioned transverse at 15 µm thickness on a Leica Microm CM1950 cryo-stat. Sections were briefly dried, then washed in 1X TBST, then blocked with 2% BSA, 1% goat serum, 0.1% Triton-X-100 in 1X PBS. Antibodies were diluted in blocking buffer and incubated overnight on sections at room temperature. The following antibodies and dilutions were used: mouse anti-PAX6 (1:25; DSHB), and rabbit anti-OLIG2 (1:300; Millipore). Primary antibody was removed, sections were washed with 1X TBST three times, then incubated for 3 hours in secondary antibodies (Invitrogen) at a 1:500 dilution. Sections were washed three times in 1X TBST, then rinsed with water, dried, and cover slips were applied with ProLong Gold mounting media with DAPI. Sections were imaged on a Leica DMi8 widefield microscope. A minimum of four embryos per genotype were analyzed.

To quantify the *Olig2* and *Pax6* expressing domains, at least three sections per embryo in the thoracic region were randomly chosen, and their area was measured using the Segmented Line Tool in ImageJ. The percentage area of each domain was calculated with respect to the entire Dorsal Ventral extent of the neural tube. The data are presented as mean ± SD, and statistical comparisons were made using Student’s t-test. GraphPad software was used for statistical analysis. p-values less than 0.05 were considered statistically significant.

For electron microscopy of neural tube primary cilia, samples were fixed in 2.5% glutaraldehyde, 2% paraformaldehyde in 0.1M cacodylate buffer containing 2 mM MgCl_2_. Following fixation, samples were dissected to expose the interior of the neural tube lumen. Dissected samples were buffer washed, post-fixed in 0.1% aqueous osmium tetroxide for 1 hour, washed with ddH_2_O, dehydrated with an ascending ethanol series, and critical point dried in liquid CO_2_ using an Autosamdri 931 (Tousimis). Dried samples were sputter-coated under planetary rotation with 15 nm iridium and imaged in a ThermoFisher Scientific Teneo scanning electron microscope at 2 kV using the Everhart-Thornley and T1 detectors.

### cAMP measurement

cAMP in whole cell lysates was measured using the Direct cAMP ELISA kit (Enzo). Cells were seeded in 6-well plates at a density of 3.5x10^5^ cells per well. The next day, cells were starved for 2 hours in serum free media followed by overnight incubation in low serum media to promote ciliation. On the day of treatment, cells were treated with 100 μM IBMX diluted in low serum media and scraped into lysis buffer (0.1 M HCl, Enzo). Samples were analyzed per the manufacturer’s protocol. cAMP concentrations were calculated using a 4-parameter logistic (4PL) and cAMP concentrations were normalized to the total protein concentrations, determined by a BCA protein assay (Thermo Fisher Scientific).

### Ciliary cAMP assay

IMCD3 cells were seeded at 18,000 cells/well in an 8-well chamber slide (Nunc™ Lab-Tek™ II Chamber Slide™ System) and transduced the next day with the ratiometric cilia-targeted cADDis BacMam (Montana Molecular, D0211G) per manufacturer’s recommendation. Cells were infected with 20 µl of BacMam sensor stock in a total of 250 µl of media containing 2 mM Sodium Butyrate (Molecular Montana) for 30 minutes at room temperature in dark followed by 5 hours at 37°C. BacMam was removed and replaced with low serum media containing 1 mM Sodium Butyrate for 16-24 hours with or without vehicle, GIRI, or L-161,982 treatment. Prior to imaging, cells were incubated in PBS for 20 minutes at room temperature. Positive agonist FSK (100 µM) was added 30 seconds after the recording was initiated and images were acquired on a Nikon A1R confocal on live cell imaging mode (63x, epifluorescence) every 10 seconds for 8 minutes. For experiments with SAG, the signal was recorded for basal conditions followed by incubation with forskolin for 1.5 minutes and then treated with either SAG or vehicle and recorded every 10 seconds for 8 minutes.

For knockdown experiments, IMCD3 cells were seeded in 8-well chamber slides (Nunc™ Lab Tek™ II Chamber Slide™ System) followed by siRNA treatment the next day with either the control siRNA or *Adcy3, Adcy5*, *Adcy6* or *Ankmy2* siRNA. Cells were then transduced the following day with the ratiometric cilia-targeted cADDis BacMam and the experiment was carried out as described above.

### Generation of optogenetic bPAC cells

IMCD3 cell lines stably expressing Cyto-bPAC-GFP and Cilia-bPAC-GFP were generated using the Flp-In system following manufacturer’s instructions (Invitrogen). IMCD3 Flp-In cells (Breslow and Nachury, 2015) were transfected using Lipofectamine 3000 with the appropriate plasmids and selected with 40 mg/mL hygromycin (Gibco). Single colonies were expanded, and protein expression was confirmed by fluorescence microscopy for GFP.

### Optogenetic stimulation

bPAC-expressing IMCD3 Flp-In cells were plated onto coverslips, grown to confluency, and pretreated with either vehicle or L161-982 (10 µM) in serum-free media. After 2 hours, media was changed to low serum media containing vehicle or drug and SHH-N conditioned media (100 µL/mL), and then incubated for ∼18 hours. Prior to immunofluorescence analysis, cells were transferred to a custom-made LED humidified incubator with 5% CO_2_ for continuous blue light 450 nm stimulation at ∼0.4 mW/cm^2^ for 3 hours (Zhang et al., 2019). Following this, immunofluorescence staining was performed as described above.

### Quantitative Reverse Transcriptase Polymerase Chain Reaction (qRT-PCR)

Total RNA was extracted from cells using the RNeasy Mini Kit (Qiagen) according to manufacturer’s protocol. One thousand nanograms of RNA was used to synthesize complementary DNA (cDNA) using High-Capacity cDNA Reverse Transcription Kit (Applied Biosystems). qRT-PCR reactions were performed on a QuantStudio 7 Flex PCR machine using PowerUp Sybr Green Master Mix (Applied Biosystems). Corresponding changes in the expression levels of selected genes were calculated using the ΔΔCt method relative to housekeeping genes, *Ppia* and *Btf3* (Arensdorf et al., 2017).

### Cell Line Generation

EP_4_ knockout cells were generated from *Ep4^-/-^*embryos (Segi et al., 1998). Embryos (13.5 dpc) were freshly isolated and placed in individual 10 cm culture dishes where they were eviscerated. Heads were removed and used for genotyping to identify homozygous null embryos and wild type littermates using conventional PCR. Embryo bodies were transferred to 15 mL conical tubes containing 5 mL fresh PBS and dissociated by aspirating through a 16G needle into a 10 mL syringe and expelling the contents. Warm media (DMEM+10%FBS+1%pen/strep) was then added to each conical tube to bring the total volume to 10 mL. This solution was transferred to a 15 cm culture dish containing 25 mL additional media and placed in a 37°C incubator. After 24h, media and non-adherent tissue pieces were removed, and fresh media was added. Cells were grown to 80-90% confluence, after which they were passaged and frozen down after passage number two. *Abcc4^-/-^* NIH-3T3 cells were generated as previously described (Wijaya et al., 2020).

### PGE_2_ ELISA

IMCD3 cells were cultured in a 12-well culture plate and were pre-treated for 2 hours with either Cyclopamine (10 µM) or Giripladib (GIRI, 4 µM) in serum free medium. Following pretreatment, medium was changed to low serum containing either vehicle, Cyclopamine, or GIRI in control or SHH-N conditioned media, and then incubated for 36 hours. PGE_2_ levels were determined using by ELISA (R&D Systems) according to the manufacturer’s instructions. The absorbance of each well was measured at 450 nm with correction at 540 nm.

### siRNA Transfection

Transfections using Lipofectamine RNAiMAX were performed following the manufacturer’s protocols. Briefly, IMCD3 or NIH-3T3 cells were plated in a 6-well tissue culture plate at a density of 175,000 or 100,000 cells per well. The following day, cells were treated with 25 pmol of either the scrambled siRNA or *Adcy3, Adcy5*, *Adcy6* or *Ankmy2* siRNA. Cells were analyzed 48 hours post transfection.

### IFT88 Velocity

To analyze intraciliary trafficking, Kymographs for anterograde IFT88 velocities were generated in IMCD3 cells as previously described (Besschetnova et al., 2009). Briefly, 120,000 cells were grown on a coverslip in a 24 well plate. Cells were transfected with IFT88-GFP and 5HT6 mCherry plasmids the following day. Cells were pretreated with the EP4 inhibitor L161,982 for 2 hours and then switched to low serum media containing L161,982 either in the presence or absence of SHH-N conditioned media and incubated overnight. On the day of the experiment, the coverslips were flipped onto a 2-well chamber slide (Nunc^TM^ Lab-Tek^TM^ II Chamber Slide^TM^ System) and the cells were imaged every 200 ms for 30 s on a Nikon A1R confocal microscope with a 60x oil objective and 37 °C, 5% CO_2_ heat stage (Live Cell Instrument). The resulting videos were analyzed by ImageJ macros KymographClear and KymographDirect to generate violin plots (Mangeol et al., 2016).

## QUANTIFICATION AND STATISTICAL ANALYSIS

Quantification of ciliary lengths and SMO ciliary accumulation values were calculated from three independent experiments with at least 100 cilia per condition per experiment. A maximum fluorescence intensity projection from a sum of at least 10 slices was generated before length and ciliary intensity quantifications. SMO ciliary intensity was normalized to acetylated α-tubulin intensity. At least three randomly chosen fields of view were selected per condition per experiment, and all cilia within the field were quantified using LAS X. For ciliary SMO signal intensity determination, each cilium surface area was traced from the base to the tip using spline profile function. The average fluorescence measurement adjacent to the cilium was subtracted from ciliary fluorescence as background. The intensity values along the cilium were exported and analyzed via GraphPad Prism. Colocalization of AC3 and AC6 within the primary cilium marker ARL13B was carried out using the ImageJ plugin JACoP. Manders’ coefficients (M1 and M2) estimate the co-occurrence fraction of a fluorescent signal on one channel (ARL13B) with a fluorescent signal of another channel (AC3/AC5/6). Manders’ coefficients range from 0 to 1. The former corresponds to non-overlapping images and the latter reflects a complete co-localization between both images.

For quantification of cAMP, the red fluorescence and green fluorescence intensities over time were processed on the Nikon NIS-Elements imaging software by defining a mask for each cilium and determining the ratio of green fluorescence to red fluorescence. The data were normalized to the initial time point for each experiment (set to 100%), and the inverse of each value was plotted using GraphPad Prism.

Statistical analyses were performed using Prism 8 (GraphPad Software). Results are presented as mean ± standard deviation. For comparisons between two groups, a two tailed unpaired t test was used unless otherwise specified. For multiple group comparisons, either one-way or two-way (depending on the number of variables) ANOVA followed by multiple comparison post hoc testing was performed as indicated using Prism 8. Significance is depicted as *p < 0.05, **p < 0.01, ***p < 0.001, ****p < 0.0001, ns = not significant.

## ACKNOWLEDGEMENTS

We thank M. Hatley and members of the Ogden lab for reagents, advice, thoughtful discussion, and comments on the manuscript. We thank S. Narumiya (Kyoto University) and the Center for Animal Resources and Development (Kumamoto University) for providing *Ptger4^-/-^* frozen sperm used to generate the *Ep ^-/-^* mouse line. J. Reiter (UCSF) provided bPAC expression vectors, S. Mukhopadhyay (UTSW) provided anti-GPR161, and M. Nachury (UCSF) provided Flp-In IMCD3 cells. We thank J. Temirov, J. Messing, A. Carisey, and the St. Jude CTIC for assistance with confocal and electron microscopy. Diagrams were created with BioRender.com.

## FUNDING

This work was supported by National Institute of Health grants R35GM122546 (S.K.O.) and R01HD099777 (E.L.S.) and by ALSAC of St. Jude Children’s Research Hospital. The CTIC is supported by SJCRH and NCI P30 CA021765. The content is solely the responsibility of the authors and does not necessarily represent the official views of the funding agencies.

## AUTHOR CONTRIBUTIONS

Conceptualization, S.S.A. and S.K.O.; Methodology, S.S.A., M.A.A., B.M.Y., M.E.D., Y.Z., E.L.S., A.J., C.G.R., and S.K.O.; Validation, S.S.A., M.A.A., M.E.D., and Y.Z.; Analysis, S.S.A., M.A.A., M.E.D., Y.Z., and A.J.; Investigation, S.S.A., M.A.A., M.E.D., and Y.Z.; Resources, J.D.S., Z.R., C.G.R., N.B., E.L.S., and S.K.O.; Writing – Original Draft, S.S.A. and S.K.O.; Writing – Review & Editing, S.S.A., M.E.D., J.D.S., E.L.S., and S.K.O.; Visualization, S.S.A.; Supervision, S.K.O. All authors approved the submitted manuscript.

## DECLARATION OF INTERESTS

The authors declare no competing interests.

## SUPPLEMENTARY FIGURE LEGENDS

**Supplementary Figure 1:**
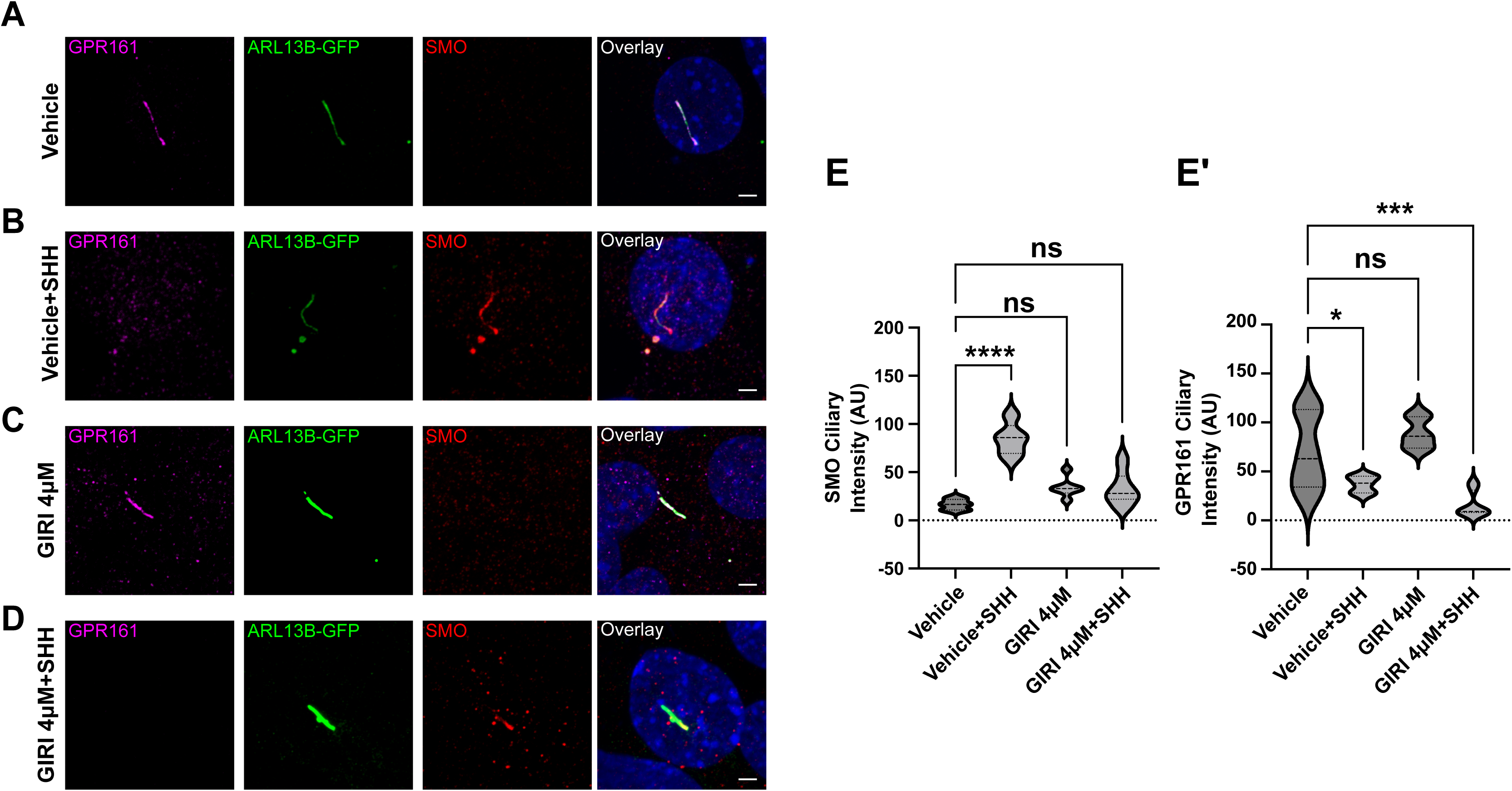
cPLA_2_ is required for SMO ciliary accumulation but not for GPR161 ciliary exit. **(A-D).** IMCD3 cells were treated with vehicle **(A-B)** or GIRI **(C-D**, 4 μM) in the absence or presence of SHH as indicated. GPR161 is magenta, SMO is red, ARL13B (green) marks primary cilia. Scale bar = 3 μm. **(E-E’)**. Quantification of ciliary signal intensity for SMO **(E)** and GPR161 **(E’)** in cells treated with SHH, GIRI, or vehicle control. The experiment was performed twice with at least 30 cilia imaged per condition per experiment and all data were pooled. Significance was determined using a one-way ANOVA. Significance is indicated as follows: *<0.05, **<0.01, ***<0.001, ****<0.0001, and ns, p > 0.05.

**Supplementary Figure 2:**
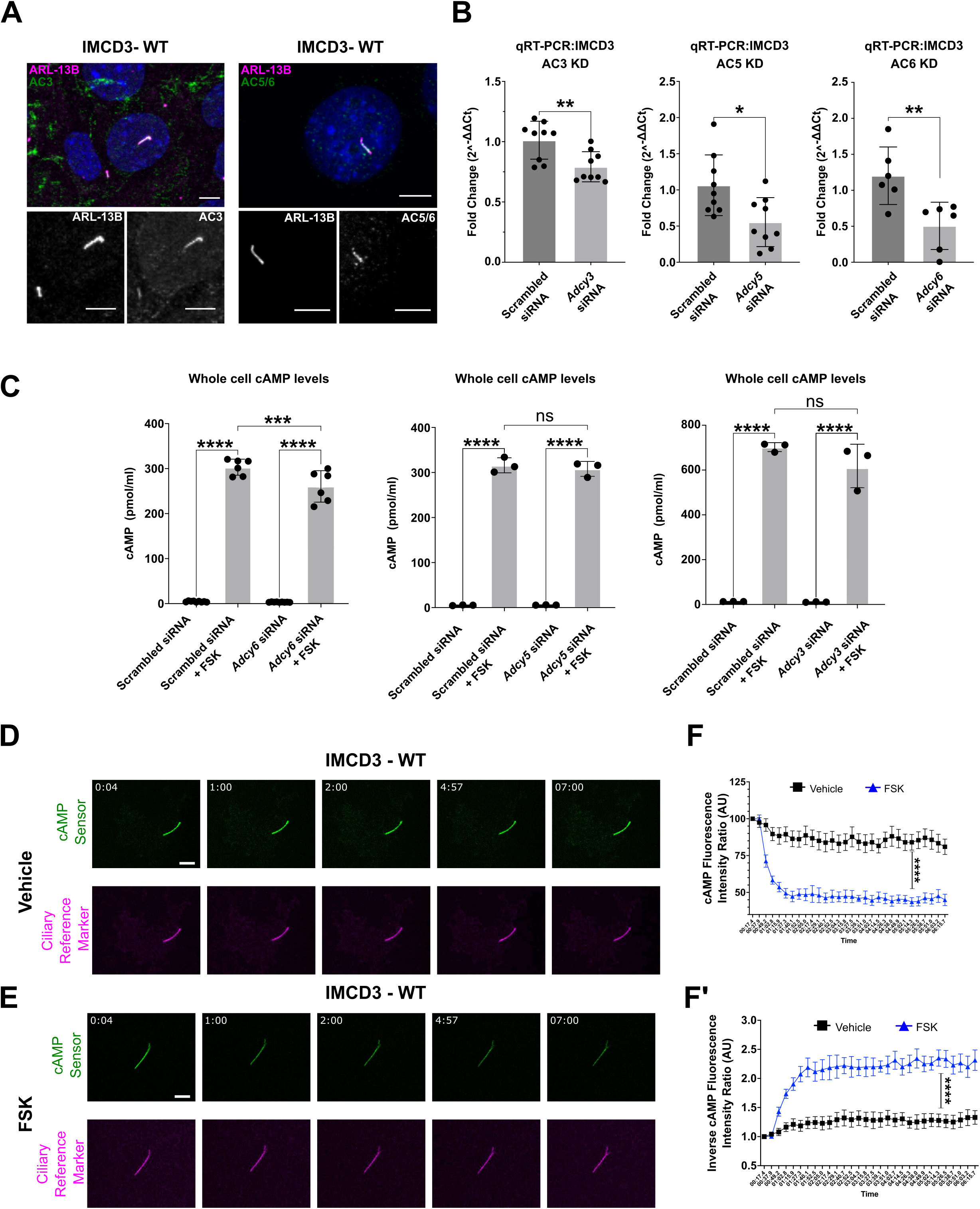
Ciliary AC controls primary cilium cAMP. **(A).** AC3 and AC5/6 localize to primary cilia in IMCD3 cells. ARL13B (magenta in merge and white in insets) marks primary cilia. Ciliary ACs are shown in green (merge) and in white (insets). Scale bar = 5μm. **(B).** *Adcy3*, *Adcy5,* or *Adcy6* knockdown in IMCD3 cells was validated by qRT-PCR ∼48 hours after treatment with AC-specific siRNA or scrambled control. Fold-change in expression was determined using the 2^−ΔΔCt^ method. Experiments were performed 3 times in triplicate and all data pooled. Statistical significance was calculated using Student’s t-test. **(C).** Knockdown of ciliary AC does not alter forskolin (FSK)-stimulated whole cell cAMP elevation. The indicated ciliary AC were knocked down as above. Cells were stimulated with 100 μM forskolin ∼48 hours after siRNA treatment and whole cell cAMP was measured using the direct cAMP ELISA assay. Assays were performed at least 3 times and all data pooled. Statistical significance was calculated using Student’s t-test. **(D-F’)**. Live imaging of cADDis-expressing IMCD3 cells shows changes in fluorescence intensity of a ciliary cAMP sensor (green) and stable fluorescence of the ciliary reference marker (magenta) following exposure to vehicle **(D)** or forskolin (FSK, 100 µM) **(E)**. Increasing cAMP decreases green fluorescence. Scale bar = 2 μm. **(F-F’).** Relative cAMP shift is shown as native **(F)** or inverse **(F’)** of the fluorescent intensity ratio of the cAMP sensor to reference marker. Inverse ratios are shown in the main figures to clearly illustrate cAMP increase. Significance was determined by calculating the area under the curve followed by Student’s t-test analysis. Significance is indicated as follows: *<0.05, **<0.01, ***<0.001, ****<0.0001, and ns, p > 0.05.

**Supplementary Figure 3:**
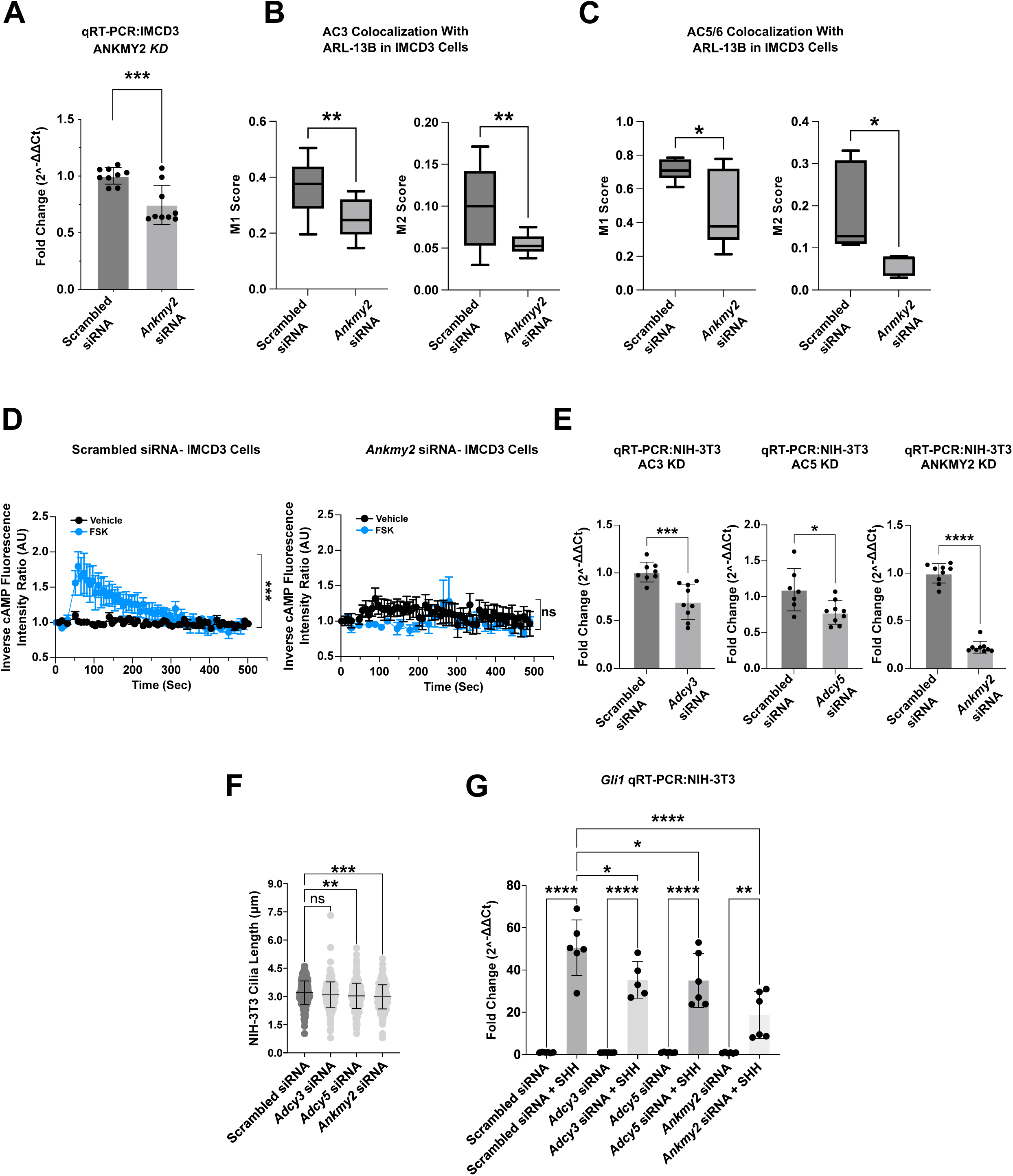
*Ankmy2* knockdown impacts IMCD3 ciliary AC enrichment and SHH-stimulated transcriptional activation. **(A).** qRT-PCR verification of *Ankmy2* knockdown in IMCD3 cells ∼48 hours after treatment with *Ankmy2* siRNA or scrambled control. Fold-change in expression was determined using the 2^−ΔΔCt^ method. **(B-C)**. Knockdown of the ciliary AC chaperone *Ankmy2* reduces primary cilium AC occupancy. Colocalization of AC3 **(B)** and AC5/6 **(C)** to ARL13B in primary cilia of IMCD3 cells in scrambled and *Ankmy2* siRNA-treated cells is shown. Both M1 and M2 scores of AC3 and AC5/6 to ARL13B are significantly reduced in cilia of the *Ankmy2* siRNA treated cells compared to control. Statistical significance was calculated using Student’s t-test. **(D)**. *Ankmy2* knockdown reduces forskolin-induced ciliary cAMP increase. Forskolin (FSK) induced changes in primary cilium cAMP were monitored in *Ankmy2* knockdown cells using the CADDis sensor. Forskolin-mediated ciliary cAMP accumulation is blocked by *Ankmy2* knockdown. Significance was determined by calculating the area under the curve followed by Student’s t-test analysis. **(E)**. *Adcy3*, *Adcy5,* and *Anmky2* knockdown in NIH 3T3 cells was validated by qRT-PCR ∼48 hours after treatment with *Adcy3, Adcy5,* or *Ankmy2* siRNA or scrambled control. Fold-change in expression was determined using the 2^−ΔΔCt^ method. Student’s t-test was used to calculate statistical significance. **(F-G)**. Knockdown of *Adcy3, Adcy5,* or *Ankmy2* shortens primary cilia and reduces SHH target gene induction in NIH 3T3 cells. **(F)**. Average lengths of ARL13B marked primary cilia of NIH-3T3 cells were determined as described above. Knockdown experiments were repeated three times with 75-100 primary cilia measured per condition per experiment. **(G)**. *Gli1* fold change was determined in control or SHH-stimulated NIH-3T3 cells following *Adcy3, Adcy5,* or *Ankmy2* knockdown. Significance was determined by one-way ANOVA. For all experiments, significance is denoted as follows: *<0.05, **<0.01, ***<0.001, ****<0.0001, and ns, p > 0.05. Data are represented as mean ± SD. qPCR analyses were performed twice in triplicate and all data were pooled.

**Supplementary Figure 4:**
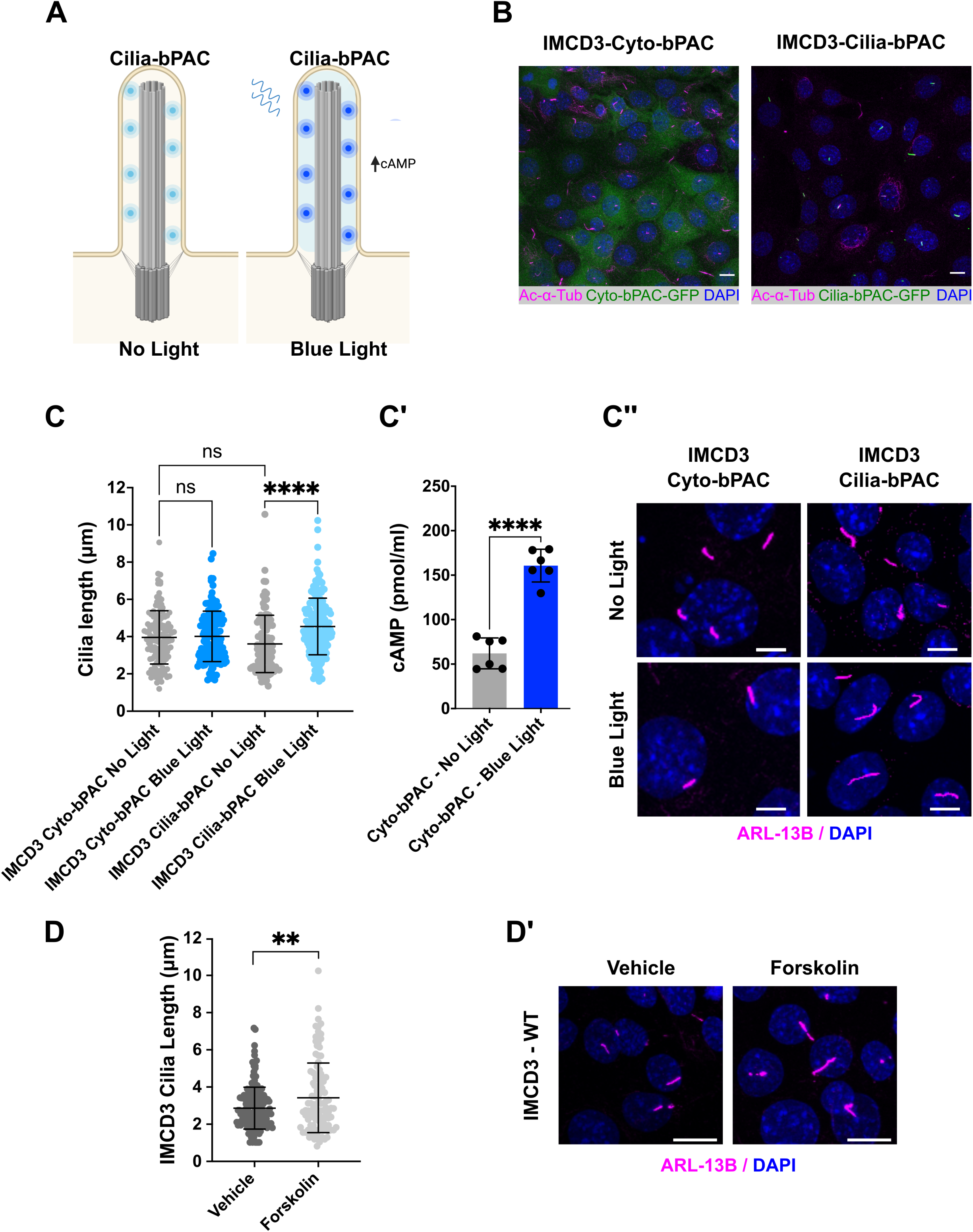
Validation of optogenetic cAMP modulation in IMCD3 cells. **(A).** Schematic of optogenetic cAMP modulation. Ciliary bPAC is activated by blue light to increase ciliary cAMP**. (B).** Immunofluorescence imaging of IMCD3 cells expressing Cyto-bPAC-GFP or Cilia-bPAC-GFP (green). Acetylated α-tubulin marks cilia (magenta) and DAPI (blue) marks nuclei. Scale bars = 10 μm. **(C-C”).** Average ciliary length was quantified in IMCD3 Cyto-bPAC, and IMCD3 Cilia-bPAC cells in control conditions or following 180 minutes of blue light exposure. **(C’).** Total cellular cAMP was measured as in Supplementary Figure 2C. Error bars indicate SD. Significance was determined by a one-way ANOVA in **(C)** and a Student’s t-test in **(C’)**. **(D-D’).** Ciliary lengths were measured in vehicle and forskolin treated cells. Average primary cilium length was calculated by measuring cilia of ≥100 cells/condition across three independent experiments. All data were pooled. **(D’).** IMCD3 cells were treated with vehicle or 100 µM forskolin. The ciliary axoneme is marked by ARL13B (magenta) and DAPI (blue) marks nuclei. Scale bar = 5 μm. Significance was determined using a one-way ANOVA. For all experiments significance is indicated as follows: *<0.05, **<0.01, ***<0.001, ****<0.0001, and ns, p > 0.05.

## Notes

### Competing Interest Statement

The authors have declared no competing interest.

### Summary of Updates

Additional controls have been added throughout the manuscript. Text has been edited for clarity.

